# Mosaic Ends Tagmentation (METa) assembly for extremely efficient construction of functional metagenomic libraries

**DOI:** 10.1101/2021.02.01.429292

**Authors:** Terence S. Crofts, Alexander G. McFarland, Erica M. Hartmann

## Abstract

Functional metagenomic libraries, physical bacterial libraries which allow the high-throughput capture and expression of microbiome genes, have been instrumental in the sequence-naïve and cultivation-independent discovery of novel genes from microbial communities. Preparation of these libraries is limited by their high DNA input requirement and their low cloning efficiency. Here, we describe a new method, METa assembly, for extremely efficient functional metagenomic library preparation. We apply tagmentation to metagenomic DNA from soil and gut microbiomes to prepare DNA inserts for high-throughput cloning into functional metagenomic libraries. The presence of mosaic end sequences in the resulting DNA fragments synergizes with homology-based assembly cloning to result in a 300-fold increase in library size compared to traditional blunt cloning based protocols. Compared to published libraries prepared by state-of-the-art protocols we show that METa assembly is on average 23- to 270-fold more efficient and can be effectively used to prepare gigabase-sized libraries with as little as 200 ng of input DNA. We demonstrate the utility of METa assembly to capture novel genes based on their function by discovering novel aminoglycoside (26% amino acid identity) and colistin (36% amino acid identity) resistance genes in soil and goose gut microbiomes. METa assembly provides a streamlined, flexible, and efficient method for preparing functional metagenomic libraries, enabling new avenues of genetic and biochemical research into low biomass or scarce microbiomes.

**IMPORTANCE:** Medically and industrially important genes can be recovered from microbial communities by high-throughput sequencing but are limited to previously sequenced genes and their relatives. Cloning a metagenome *en masse* into an expression host to produce a functional metagenomic library is a sequence-naïve and cultivation-independent method to discover novel genes. This directly connects genes to functions, but the process of preparing these libraries is DNA greedy and inefficient. Here we describe a library preparation method that is an order of magnitude more efficient and less DNA greedy. This method is consistently efficient across libraries prepared from cultures, a soil microbiome, and from a goose fecal microbiome and allowed us to discover novel antibiotic resistance genes. This new library preparation method will potentially allow for the functional metagenomic exploration of microbiomes that were previously off limits due to their rarity or low microbial biomass, such biomedical swabs or exotic samples.

## INTRODUCTION

The widespread adoption of high-throughput DNA sequencing technology has resulted in a new and deserved appreciation for the genetic diversity present in microbial communities aka microbiomes (1). Projects studying the chemical biology of microbiomes have shown that this genetic diversity translates into enormous biochemical diversity (2–6). However, linking genes from microbial community genetic material (the metagenome) to biochemical activity remains difficult due to current limitations in gene prediction and annotation of specific activities. Direct observation of biochemical function *in vitro* or phenotype *in vivo* remain the gold standards of functional assignment as a result (7, 8).

One method that unites the culture- and sequence-independence of high throughput sequencing with the functional observations that result from cloning and expression studies is functional metagenomics. Functional metagenomics relies upon the construction of metagenomic libraries in which a portion of a microbiome’s metagenome is captured in a bacterial artificial chromosome (BAC) or plasmid library and housed in an expression host, often *E. coli* (2) (**Figure 1**). This technique allows function to be linked directly to genes without requiring laboratory growth of the originating organisms or prior knowledge of target gene sequence. Functional metagenomic libraries have been used to bioprospect for novel bioactive compounds ((9) and reviewed in (3, 10)), novel enzymes of potential interest to industry (11, 12), and enzymes useful in the production of biofuels (13, 14) (see also reviews (15–17)) and have been created using metagenomic DNA from environments as varied as soils, adult and infant fecal samples, sewage and waste-water effluent, and animal samples (18, 19, 28–35, 20–27). One particularly successful application has been the identification of antimicrobial resistance genes (7, 36) that would not have been identified by sequencing due to their low predicted amino acid identity.

**Figure 1.**
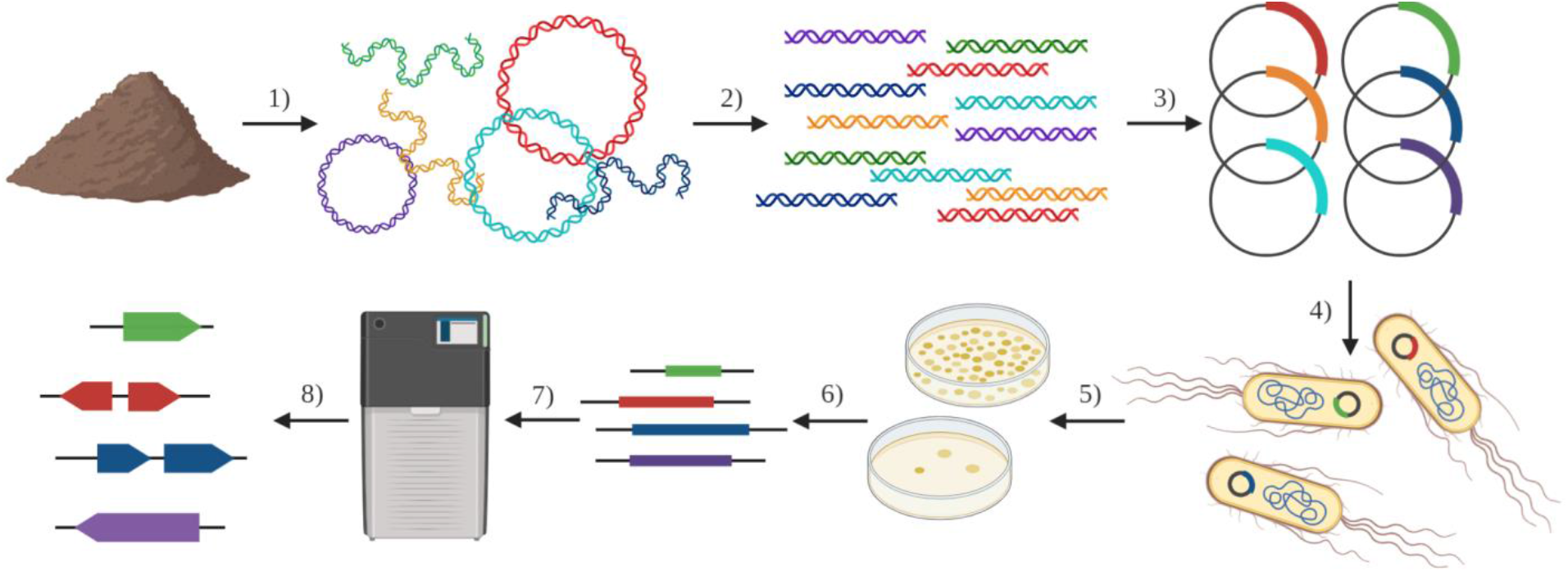
Functional metagenomic library pipeline. The general pipeline for the creation and use of functional metagenomic libraries to capture and discover genes from metagenomes. 1) Extraction of metagenomic DNA from a microbiome (*e.g.* soil or fecal samples). 2) Fragmentation of metagenomic DNA to desired size range (*e.g.* by sonication, restriction enzyme digestion, or tagmentation). 3) Cloning of fragments into expression vectors following size selection (*e.g.* by blunt ligation or homology-based assembly. 4) Transformation *en masse* of vectors into an expression host (*e.g. E. coli* DH10B) to create functional metagenomic library. 5) Functional selection or screen of library (*e.g.* on antibiotics to select for resistance). 6) Amplification of selected inserts using vector-specific primers. 7) High-throughput sequencing of selected metagenomic amplicons (*e.g.* by Illumina or PacBio technologies). 8) Annotation of sequenced amplicons to link novel genes with selected/screened function (*e.g.* discovery of novel aminoglycoside acetyltransferases). Figure created in BioRender.com.

The basic steps for creating a functional metagenomic library consist of metagenomic DNA extraction, DNA fragmentation, cloning of fragments into a plasmid/BAC/*etc*., and transformation of the plasmid library into an expression host (6, 36) (**Figure 1**). In many cases, the libraries used in these experiments contain gene-sized inserts (*ca.* 1 kb to 10 kb) and make use of sonication or other physical means to fragment metagenomic DNA into inserts followed by blunt cloning into an expression vector (18, 21, 25, 26, 33, 37). These two steps, sonication and blunt cloning (steps 2 and 3 in **Figure 1**), greatly lower the potential efficiency of functional metagenomic library creation, necessitating multiple micrograms of input DNA mass.

Like functional metagenomic libraries, shotgun sequencing libraries have, until recently, relied largely upon physical methods for DNA fragmentation and have similarly required substantial input DNA mass. In contrast, transposase enzymatic DNA fragmentation (38, 39) (tagmentation, known commercially as Nextera) produces DNA fragments using transposase enzymes that create mostly random (40) double stranded breaks in target DNA by insertion of their mosaic end sequence oligo cargo (38). This method substantially decreases costs and input DNA mass requirements (38–43) but has not been applied yet to the preparation of functional metagenomic libraries. We hypothesized that tagmentation reactions could replace sonication in the preparation of functional metagenomic libraries (**Figure 2A**), decreasing input DNA requirements by greater than 10-fold compared to current methods (**Figure 2B**). Tagmentation also results in incorporation of 19 bp transposase oligos on the 5’ ends of both strands of fragmented DNA. We hypothesized that incorporation of these sequences on the ends of inserts could allow us to use homology-based DNA assembly protocols (*e.g.* Gibson assembly(44), *etc*) in place of blunt ligation (**Figures 2C** and **2D**). We hypothesized that incorporation of matching sequences in an expression vector (**Figure 2E**) would allow the vector to capture inserts by hybridization leading to significantly increased efficiency in library preparation.

**Figure 2.**
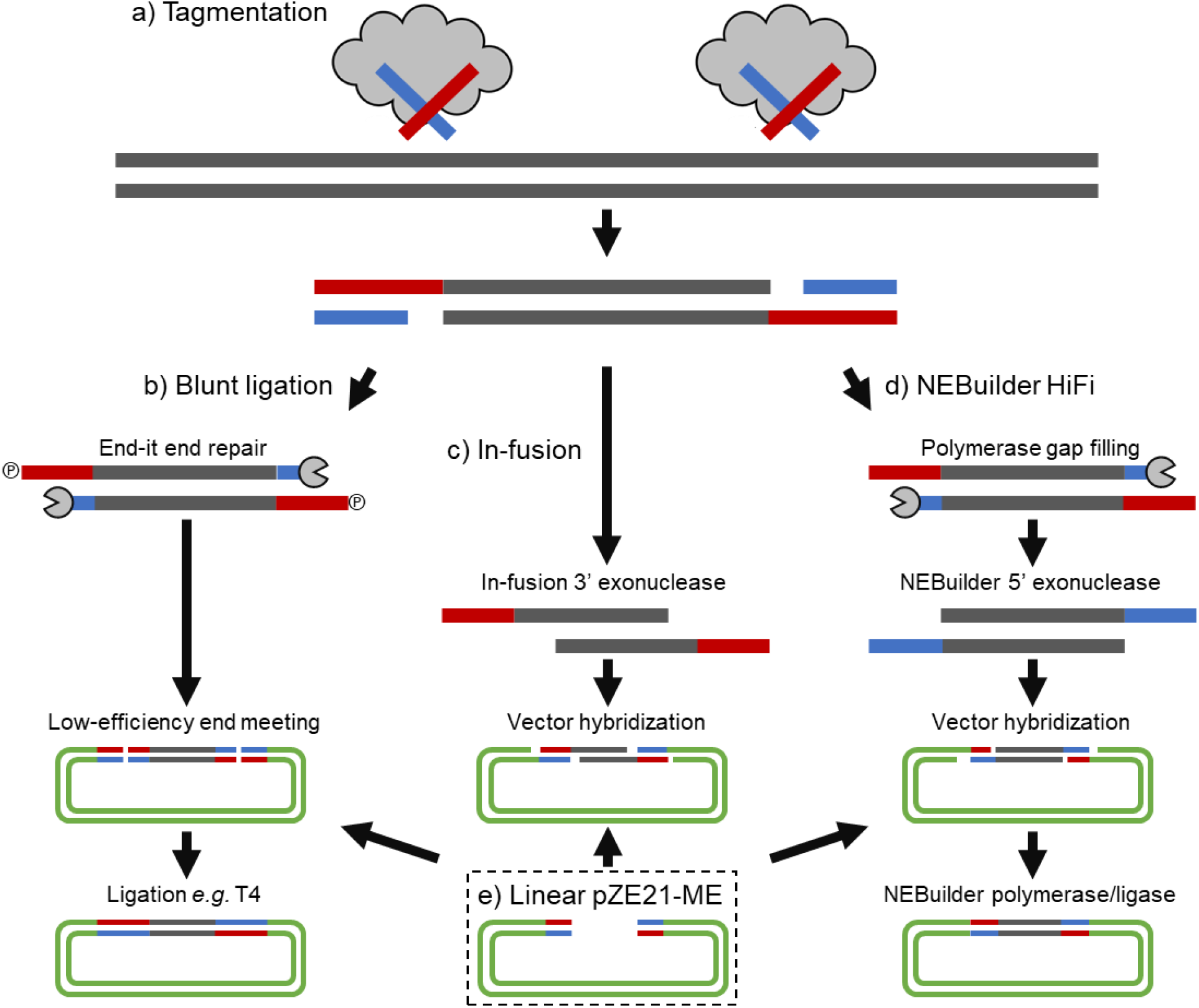
Blunt cloning protocol compared to METa assembly with NEBuilder HiFi or In-Fusion. a) Transposase enzyme fragments DNA with 5’ mosaic end oligos. Inserts can be used as input for all three methods. All three protocols are compatible with linear pZE21-ME vector prepared by inverse PCR. b) Blunt cloning *via* end-repair and ligase. 5’ overhangs must be resolved by gap filling and phosphorylation using end-repair enzyme mixes. Blunt ended inserts can be ligated to blunt ended vector. c) METa assembly *via* In-Fusion enzyme mix. In-Fusion 3’ exonuclease activity is directly compatible with transposase fragments. Single stranded DNA overhangs on inserts and vector hybridize into a stable complex that can be transformed without filling gaps or covalently sealing nicks. d) METa assembly *via* NEBuilder HiFi enzyme mix. 5’ overhangs must be resolved by DNA polymerase gap filling. NEBuilder HiFi enzyme mix includes 5’ exonuclease to create 3’ overhangs which hybridize with target pZE21-ME. DNA polymerase fills in gaps and ligase seals nicks. e) pZE21-ME is prepared and linearized by inverse PCR and is compatible with all three DNA pipelines.

Here we report our testing of these hypotheses and the development, validation, and application of a new general method for functional metagenomic library preparation that we are calling <u>M</u>osaic <u>E</u>nds <u>Ta</u>gmentation assembly, or METa assembly. Our method takes advantage of the unexpected synergy between tagmentation and assembly cloning to produce functional metagenomic libraries with up to 270-fold more efficiency and 25-fold reduced input DNA mass requirements compared to the current method for functional metagenomic library preparation. METa assembly has the potential to greatly improve and expand the field bioprospecting, catalyzing the discovery of novel microbial chemistry from genetically diverse microbiomes.

## MATERIALS AND METHODS

### Preparation and purification of transposase enzyme

Expression and purification of transposase enzyme was carried out based on modifications of protocols published by Picelli *et al.* and Hennig *et al.* (41, 42). *E. coli* XL1 blue carrying the pTXB1-Tn5 plasmid was a gift from Rickard Sandberg (Addgene plasmid #60240; http://n2t.net/addgene:60240; RRID:Addgene_60240) and maintained as specified. The pTXB1-Tn5 plasmid was recovered from an *E. coli* culture grown in LB supplemented with 100 μg/ml carbenicillin (LB+CA100) *via* miniprep kit (New England Biolabs, cat#T1010S). The plasmid was transformed into chemically competent *E. coli* BL21(DE3) cells (New England Biolabs, cat#C2527I) following manufacturer recommendations and selected on LB+CA100 before maintenance as a 15% glycerol stock at −80°C. A single colony of this strain was used to inoculate 1 ml of LB+CA100 and incubated shaking overnight at 37°C. In the morning, the saturated overnight culture was used to inoculate 1 L of Studier ZYM-5052 auto-induction media (45) in a 2.8 L Fernbach flask. The culture was grown aerobically at 37°C until it began to turn turbid by eye, approximately 3 hours, at which point the temperature was decreased to 20°C and shaking maintained at 350 rpm overnight. Cells were collected by centrifugation at 8,000 rcf for 20 min at 4°C after OD_600_ measurements suggested growth had plateaued at an OD_600_ of ∼3.72 AU. The resulting wet cell pellet weighed 8.08 g and a sample was analyzed by SDS-PAGE (sodium dodecyl sulfate polyacrylamide gel electrophoresis) to verify induction of the ∼75 kDa Tn5-chitin binding domain fusion protein.

The cell pellet was resuspended to 10% w/v in HEGX buffer composed of 20 mM HEPES buffer pH 7.2, 0.8 M NaCl, 1 mM ethylenediaminetetraacetic acid (EDTA), 10% v/v glycerol, and 0.2% v/v triton X-100 and supplemented with 20 μM phenylmethylsulfonyl fluoride (PMSF) as a protease inhibitor. Resuspended cells were lysed on ice by sonication using a W-225 sonicator with 6 cycles of 1 min on and 1 min off at output 5, 50% duty. Insoluble debris was removed by centrifugation at 15,000 rcf for 30 min at 4°C following which 2.1 ml of neutralized 10% polyethyleneimine (Millipore Sigma, cat#P3143) was added to the decanted supernatant dropwise while stirring at 4°C. The precipitated *E. coli* genomic DNA was removed by centrifugation for 10 min at 9,000 rcf at 4°C.

The fusion protein was purified from the clarified supernatants by adding 20 ml of chitin resin (New England Biolabs, cat#S6651S) and incubating with gentle rotation overnight at 4°C. The resin was washed with approximately 400 ml of HEGX buffer, following which the drained resin was added to 30 ml of HEGX buffer supplemented with 100 mM β-mercaptoethanol to initiate intein cleavage of the Tn5 transposase from the chitin binding domain. Cleavage was allowed to proceed with gentle rotation at 4°C for approximately 48 hr after which transposase was collected from the resin by draining and saving the flow-through. The resin was washed with 2X dialysis buffer consisting of 100 mM HEPES pH 7.2, 0.2 M NaCl, 0.2 mM EDTA, 20% w/v glycerol, 0.2% triton X-100, and 2 mM dithiothreitol (DTT) and washes were pooled with the initial elution. The pooled eluates were concentrated and exchanged into 2X dialysis buffer using an Amicon Ultra 15 ml 3,000 molecular weight cut-off filter (Millipore Sigma, cat#UFC900324). This concentrate contained 1.85 mg/ml protein as determined by BCA assay (Thermo Scientific, cat#23225) in *ca.* 5.67 ml 2X dialysis buffer. After adding 6.2 ml of glycerol and 1.89 ml 2X dialysis buffer enzyme stocks were stored at −20°C as 500 μl aliquots. Final transposase stocks contained approximately 763 ng/μl protein.

### Verification of transposase activity

Enzyme activity was verified by observing degradation of a soil metagenomic DNA extract. Mosaic end primers 5Phos_METagA1 and METagA2 (**Supplemental table 1**) synthesized by Integrated DNA Technologies (IDT Inc.) were brought to a concentration of 100 μM in 50 mM NaCl, 40 mM Tris pH 8. Annealing was carried out by combining 10 μl aliquots of each oligo and incubating in a thermocycler using the following settings: 5:00 at 95°C, cool to 65°C at 0.1°C/sec, hold at 65°C for 5:00, and cool to 4°C at 0.1°C/sec. Annealed oligos were maintained as aliquots at −20°C until use. Annealed mosaic ends oligos were loaded into transposases by adding 0.143 volumes of annealed oligos to one volume of 763 ng/µl transposase stock and incubating at room temperature (*ca.* 23°C) for 1 hr.

As a test substrate, metagenomic DNA was extracted from a soil sample taken from the Northwestern University campus (coordinates 42.05662, −87.674247) using a DNeasy PowerSoil Kit (Qiagen, cat#12888-100). Quantification of DNA for all experiments was performed using a QuantiFluor ONE dsDNA system (Promega, cat#E4871). Test tagmentation reactions were performed by combining MilliQ water (to 20 μl final volume), 4 μl of 5X TAPS-DMF buffer (50 mM TAPS buffer pH 8.5, 25 mM MgCl_2_, 50% v/v dimethylformamide), 1 μl of 50 ng/μl soil metagenomic DNA, and 1 μl of water (control) or 1 μl of loaded transposase (corresponding to 665 ng). Reactions were incubated in a thermocycler for 7 min at 55°C at which point reactions were quenched by addition of 5 μl of 0.2% SDS (final concentration 0.05% SDS) and incubation at 55°C for 5 min. For each 25 μl quenched reaction (+/- transposase), 12.5 μl was purified using a silica column-based kit (New England Biolabs, cat#T1030S, polymerase chain reaction (PCR) purification kit used for all future enzymatic clean-up steps) and eluted with 12.5 μl water while the remaining 12.5 μl were not purified further. Next, 2.5 μl of 6X loading dye was added to both the un-purified reactions and purified reactions and samples were analyzed on an agarose gel. Unless otherwise noted, all agarose gel experiments were conducted using ∼0.7% agarose pre-cast with SYBRSafe (Invitrogen, cat#S33102) following manufacturer’s recommendations. Conversion of the high molecular weight DNA smear into a low molecular weight smear confirmed active transposase.

### Tagmentation of high molecular weight DNA to produce 1 kb to 10 kb fragments

High molecular weight DNA for use as a test transposase substrate was obtained from cultures of *Pseudomonas* sp. Strain PE-S1G-1 (referred to as ABC07) and *Pandoraea* sp. Strain PE-S2T-3 (referred to as ABC10) (46–48). Each strain was grown in 1 ml of LB+CA100 at 30°C for 48 hours and genomic DNA was extracted using a Quick-DNA High Molecular Weight kit (Zymo Research, cat#D6060) according to the manufacturer’s protocol, quantified, and combined to give a 100 ng/μl stock solution with equal input by mass from each genome.

Tagmentation reactions were assembled in 20 μl volumes in 1X TAPS-DMF buffer containing 200 ng of DNA and assembled transposase in concentrations ranging from 0 ng enzyme per ng DNA to 2 ng enzyme per ng DNA. Following incubation and quenching (see above) 6X loading dye was added to each reaction and samples were analyzed by pulsed field gel electrophoresis at 4°C using a Pippin Pulse power supply (Sage Science, cat#PPI0200) running pre-set protocol #4: 16 hr, 75 V on a 0.75% TAE agarose gel. Following electrophoresis, the gel was stained with SYBRSafe and visualized under UV light.

### Preparation of pZE21-ME vector

The pZE21 plasmid (49) was used as a template for an inverse PCR reaction to replace the multiple cloning site with tandem mosaic end sequences of AGATGTGTATAAGAGACAG-CTGTCTCTTATACACATCT. The construct was amplified by 2-step PCR according to manufacturer recommendations using Q5 high fidelity polymerase (New England Biolabs, cat#M0494S) with primers 6469TSC and 6470TSC (**Supplemental table 1**). The vector product was sized and purified from an agarose gel and circularized by end repair/phosphorylation (Lucigen, cat#ER0720) and blunt ended ligation (Lucigen, cat#LK0750H) following manufacturer instructions. The construct was transformed into *E. coli* DH10B (New England Biolabs 10-beta, cat#C3020K), selected on LB supplemented with 50 μg/ml kanamycin (LB+KA50), and frozen at −80°C as a 15% glycerol stock. Incorporation of the mosaic ends sites 23 bp downstream of the pZE21 ribosome binding site was confirmed by Sanger sequencing.

### Comparison of METa assembly using In-fusion or NEBuilder HiFi assembly to blunt ligation

Triplicate assembly/ligation tests were carried out to determine the feasibility of using mosaic end tags as targets for homology-mediated assembly of metagenomic cloning reactions. Soil metagenomic DNA (see above) was used as input DNA in a bulk transposase reaction containing 477.4 μl autoclaved MilliQ water, 124 μl freshly prepared 5X TAPS-DMF buffer, 15.5 μl of 290 ng/μl metagenomic DNA (7.25 ng DNA per μl final), and 3.1 μl of 665 ng/μl assembled transposase (0.46 ng transposase per ng metagenomic DNA) in 620 μl total volume. The bulk reaction was incubated at 55°C for 7 minutes and quenched by addition of 3.1 μl of 10% SDS (final concentration 0.05% SDS) followed by incubation at 55°C for 5 minutes. The bulk reaction was purified and concentrated by PCR purification kit according to manufacturer’s directions for targeting fragments >2 kb in length. Tracking dye was added to the elution and the entire sample was loaded onto an agarose gel and run at 70V for 120 minutes alongside a λ DNA BstEII digest ladder (New England Biolabs, cat#N3014S). Prior to running, the electrophoresis apparatus and gel tray were washed with milliQ water, soaked in 10% bleach for 15 minutes, and rinsed with milliQ water. The bleaching process and use of λ DNA ladder are both necessary to prevent contamination of the metagenomic DNA fragments by foreign DNA that can be mistakenly incorporated during blunt ligation cloning (37). Fragments between ∼1 kb and ∼8 kb were size selected by excision with a clean razor blade and DNA was purified from the agarose by a silica column-based gel extraction kit (New England Biolabs, cat#T1020S, used in all gel extraction steps) with ∼500 mg of agarose used per purification column. For downstream cloning reaction calculations, the average insert size was assumed to be 4 kb. Triplicate In-Fusion, NEBuilder, and blunt ligation reactions all used this pool of inserts as their DNA inputs.

In-Fusion assembly reactions were prepared in triplicate containing 175 ng of DNA fragments, 50 ng of pZE21-ME vector (a ∼2:1 insert to vector ratio), 4 μl of 5X In-Fusion premix (Takara Bio, cat#638909), and milliQ water up to 20 μl. Reactions were held at 50°C for 15 minutes then transferred to ice according to manufacturer’s recommendations.

DNA for NEBuilder HiFi assembly reactions was prepared by combining 5 μl of Q5 high fidelity polymerase previously heated to 98°C for 30 seconds with 5 μl of DNA fragments (600 ng) and incubated at 72°C for 15 minutes to fill in potential 5’ overhangs. DNA fragments were purified by silica column as before. Triplicate NEBuilder HiFi assembly reactions were prepared according to manufacturer’s protocols containing ∼2:1 insert to vector as follows: 10 µl of 2X NEBuilder HiFi master mix (New England Biolabs, cat#E2621S), 158.7 ng of DNA fragments, 45.2 ng pZE21-ME, and milliQ water to a volume of 20 μl. Reactions were incubated at 50°C for 15 minutes then transferred on to ice.

Preparation of a metagenomic DNA library by blunt ligation was based on a modified version of the Dantas lab protocol (21, 37). DNA fragments for blunt cloning were repaired using an End-It DNA End-Repair Kit (Lucigen, cat#ER0720) according to the manufacturer’s protocol, with the single 50 μl reaction containing 600 ng of DNA fragments, 5 μl each of 10X End-Repair buffer, dNTP mix, and ATP mix, and 1 μl of End-Repair Enzyme mix in milliQ water. The reaction was held at room temperature (*ca.* 23°C) for 45 minutes then purified by silica column as before. Triplicate blunt ligation reactions were prepared with each reaction containing 115 ng of end-repaired DNA, 21.9 ng of pZE21-ME (for a ∼5:1 insert to vector ratio), 5 μl of 2X Blunt/TA Ligase master mix (New England Biolabs, cat#M0367S), and milliQ water up to 10 μl total volume. Blunt ligation was carried out at room temperature (*ca.* 23°C) for 45 minutes, as suggested by the ligase manufacturer, then the reactions were transferred to ice.

All nine reactions (three techniques, each in triplicate) were de-salted and purified using silica column kits for DNA fragments >2 kb and eluted in 10 μl of 50°C milliQ water. *E. coli* DH10B cells were made electrocompetent following a modified protocol (50) and tested to have a moderate transformation efficiency of 4.7×10^8^ cfu/μg of pUC19 DNA. Nine 45 μl aliquots of homemade competent cells stored at −80°C were thawed and, sequentially, combined with 10 μl of purified DNA in a 0.1 cm Gene Pulser cuvette (Bio-Rad, cat#1652083) and electroporated on an Electroporator 2510 instrument (Eppendorf) at 1.8 kV with default settings of 10 μF capacitance and 600 Ω resistance (note, this results in a similar τ constant to standard settings on other instruments of 25 μF capacitance and 200 Ω resistance). Cells were immediately rescued with 1 ml of room temperature (*ca.* 23°C) SOC media (Takara Bio) and recovered by incubating with shaking for 1 hour at 37°C.

Following recovery, 50 μl of each transformation reaction was plated onto LB+KA50 agar at 50-fold, 500-fold, and 5,000-fold dilution (equivalent to plating 1 μl, 0.1 μl, and 0.01 μl of the 1 ml recovery sample) and incubated overnight at 37°C. Colony counts from each titer plate were taken and used to estimate the post-recovery population density as colony forming units per ml (cfu/ml). To estimate average insert size and the rate of successful insert capture, individual colonies were picked into 120 μl of milliQ water and used as template for 15 μl colony PCR as follows: 0.3 μl each of 10 μM primers 6463TSC and 6464TSC (**Supplemental table 1**), 1.5 μl of colony suspension, 7.5 μl of One*Taq* Quick-Load 2X master mix (New England Biolabs, cat#M0486), and milliQ water to 15 μl. The reactions were incubated in a thermocycler with the following program: 5:00 at 94°C followed by 25 cycles of 0:30 at 94°C, 0:45 at 62°C, and 8:00 at 68°C, followed by 5:00 at 68°C and storage at 4°C. Reactions were analyzed on agarose gel and average insert size was calculated by comparison to a DNA ladder. The proportion of colonies with an insert was estimated by taking the proportion of reactions returning a >500 bp product (500 bp being the expected amplicon size for a vector backbone only reaction) over the total number of successful reactions. The library size for each assembly or cloning reaction was calculated using the following equation (cfu is calculated from cfu/ml * 1 ml total recovery volume):

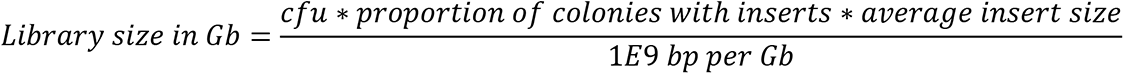

Following library size estimation, each library was normalized to the quantity of insert DNA used for the cloning step (*i.e.* In-Fusion, NEBuilder HiFi, or blunt ligation step) for direct comparison across techniques in the form of Gb library/ng insert DNA.

### Repeat comparison of NEBuilder HiFi METa assembly to blunt ligation

To confirm the efficacy of the NEBuilder HiFi METa assembly protocol, we prepared triplicate libraries by METa assembly or blunt ligation using as input mixed genomic DNA from *Pseudomonas* sp. PE-S1G-1 and *Pandoraea* sp. PE-S2T-3 (prepared above). A 1 ml tagmentation reaction consisting of 1X TAPS-DMF buffer, 10 μg mixed genomic DNA (10 ng input DNA per μl final volume), and 5 μg assembled transposase (0.5 ng of transposase per ng of input DNA) was incubated, quenched, concentrated by PCR purification kit, and size selected and purified from agarose gel as before, targeting DNA fragments between ∼1 kb and ∼8 kb.

NEBuilder HiFi assembly input DNA fragments were modified as before in triplicate reactions (PCR overhang filling by Q5 polymerase). Inserts for blunt ligation cloning were also modified in triplicate reactions (end repair by End-It kit). For both sets of triplicate reaction each replicate used 300 ng of size-selected DNA fragments as input. NEBuilder HiFi assembly was performed as before, with triplicate reactions containing 40 ng of inserts and 20 ng of pZE21-ME linear vector, and parallel triplicate sham reactions containing milliQ water in place of inserts. Blunt ligation reactions were prepared to follow established functional metagenomic library cloning protocols (21, 37). Triplicate reactions were prepared containing 1.5 μl of Fast-Link 10X ligation buffer, 0.75 μl ATP solution (10 mM), 100 ng of insert fragments, 40 ng of linearized pZE21-ME, 1 μl of Fast-Link DNA ligase (Lucigen, cat#LK0750H), and milliQ water up to 15 μl total volume. In parallel, another triplicate set of sham reactions were prepared with milliQ water replacing the corresponding volume of insert DNA. Blunt ligation reactions were incubated at room temperature (*ca.* 23°C) overnight then heat inactivated at 70°C for 15 minutes.

All 12 reactions (two techniques with triplicate insert reactions and triplicate sham reactions each) were purified and de-salted by silica column kit. For each reaction, the entire 10 μl milliQ water elution was electroporated into a 25 μl aliquot of commercial 10-beta electrocompetent *E. coli* DH10B cells (advertised transformation efficiency of 2×10^10^ cfu/μg DNA), immediately rescued in 1 ml of 37°C SOC outgrowth medium (New England Biolabs, cat#B9020S) and incubated with shaking at 37°C for 1 hour. Following recovery, 100 μl of 100-fold (10^2^), 10,000-fold (10^4^), and 1,000,000-fold (10^6^) diluted cultures were plated onto LB+KA50 plates overnight at 37°C. The remaining stocks of the sham reactions were discarded, while the remaining stocks of the insert reactions were individually used to inoculate 50 ml of LB+KA50 media and shaken O/N at 18°C before being transferred to a 37°C shaking incubator the next day to amplify the libraries to an OD_600_ of between 0.6 and 1.0 AU.

Library size was estimated as above, by counting the titer plates to find the number of colony forming units present post-electroporation and by using colony PCR to determine the proportion of cells containing an insert and to determine the average insert size. The libraries amplified in 50 ml of LB+KA50 were concentrated by centrifugation at 4,000 rcf for 7 minutes and resuspended to 10 ml in 15% glycerol in LB+KA50 media. The concentrated libraries were aliquoted 1 ml at a time into cryovials for storage at −80°C.

### Assembly of a large soil microbiome metagenomic library by METa assembly

The previously extracted and purified soil metagenomic DNA was used as input for a 5 μg tagmentation reaction performed in 1X TAPS-DMF buffer with 10 ng/μl final metagenomic DNA concentration and 0.5 ng of annealed transposase per ng of metagenomic DNA. The reaction was incubated at 55°C for 7 minutes then quenched for 5 minutes at 55°C by adding SDS to 0.05%. The reaction was purified and concentrated by silica column clean-up as before and eluted twice with 6 μl of 55°C elution buffer. To the full elution was added 1 μl of 6X tracking dye and 2 μl of glycerol before loading onto an agarose gel and running at 75V for 2 hr alongside a BstEII digest of λ DNA ladder. Fragments were size selected by excision from agarose gel, targeting inserts between 1 kb and 6 kb in length and purified by silica column and eluted twice with 25 μl elution buffer heated to 55°C. Fragments overhangs were filled by PCR with 50 μl of Q5 2X master mix for 15 min at 72°C, purified by silica column, eluted twice with 6 μl of 55°C water and quantified. Out of a total of 1.122 μg of available purified inserts two libraries were assembled. The first test library used 175 ng of inserts (∼0.14 pmol assuming 2 kb average size) assembled into 100 ng of pZE21-ME (∼0.07 pmol) using NEBuilder Hifi enzyme mix as before in a 20 μl total volume. The second library used the remaining inserts in a scaled up reaction volume of 54 μl total maintaining the recommended 0.2 pmol of total fragments per 20 μl of reaction. The assembly reactions were de-salted by silica column purification, eluted twice with 6 μl 55°C water (the 175 ng assembly) or eluted twice with 25 μl 55°C water (the bulk reaction), and electroporated into 25 μl (the 175 ng assembly) or 125 μl (the bulk assembly) of NEB 10-beta electrocompetent *E. coli* cells at 1.8 kV. The libraries were amplified overnight in LB+KA50 and stored in 1 ml glycerol stocks as before. Library sizes were quantified by colony counting and colony PCR. Colony PCR was performed using the high processivity polymerase Q5, with each 10 μl colony PCR reaction containing 0.5 μl each of primers TSC6463 and TSC6464, 5 μl of 2X Q5 polymerase, 4 μl of water. Template consisted of a sterile toothpick touched to a colony and dipped into the reaction solution. PCR reactions were carried out by heating to 98°C for 3:00, followed by 25 cycles of 98°C for 0:10 and 72°C for 4:00, followed by holding at 72°C for 5:00.

### METa assembly of soil or goose fecal metagenomic libraries using limited input DNA

A library to test the limits of METa assembly was prepared using a 10 kb DNA amplicon as input. To generate the input DNA for this assembly a 50 µl Q5 PCR reaction using template DNA and primers from a Phusion HiFi amplification control (Thermo Scientific, cat#F553S) was run as follows: 25 µl of Q5 2X hot start master mix, 6.25 µl of 4 μM primers P1 and P2 and 1 µl of template DNA (Thermo Scientific, cat#F553S), and milliQ water up to 50 μl. Amplification was carried out according to manufacturer’s protocols with a T_A_ of 65°C. The size and purity of the 10 kb product was verified by agarose gel electrophoresis, purified by silica column and used as substrate in a 200 ng, 20 μl tagmentation reaction (10 ng DNA per μl reaction, 0.5 ng assembled Tn5 per ng input DNA). Size selection, overhang filling, and purification were performed as before. Fragments (59.4 ng) were assembled into pZE21-ME vector (30 ng) by NEBuilder HiFi assembly master mix (total volume 20 μl), purified, and transformed into electrocompetent 10-beta *E. coli* cells to produce a library for quantitation by colony counting and colony PCR.

A soil functional metagenomic library was prepared as above, with 250 ng of previously extracted soil metagenomic DNA used as input in a 25 μl tagmentation reaction. The reaction was purified by silica column and fragments between 2 kb and 6 kb in length were purified by extraction from agarose gel. Overhang filling, purification and assembly by NEBuilder HiFi master mix (35.2 ng of inserts, 21 ng of linearized pZE21-ME, 20 μl total volume) were performed as before followed by assembly purification and transformation into electrocompetent 10-beta *E.* coli. The library was quantified by colony counting and colony PCR and amplified overnight as above followed by storage at −80°C in 10 x 1 ml aliquots.

To prepare a goose fecal microbiome library a freshly voided adult Canada goose (*Branta canadensis*) (**Supplemental figure 11A**) fecal pellet was collected from the Northwestern University campus (coordinates 42.056084, −87.670661). Fecal pellet collection was approved by the Northwestern University Institutional Animal Care and Use Committee (IACUC) under protocol EC20-0252. Within 15 minutes of collection, 474 mg of fecal pellet was used as input for metagenomic DNA extraction using a DNeasy PowerSoil Kit (Qiagen, cat#12888-100) following modifications for goose microbiome extraction suggested by *Cao et. al* 2020 (51). Briefly, these consisted of incubating the sample suspended in Qiagen buffer CD1 at 65°C for 10 minutes followed by incubation at 95°C for 10 min followed by the kit manufacturer’s protocol.

Tagmentation was performed as above (10 ng input DNA per μl volume, 0.5 ng of assembled transposase per ng of input DNA) using 300 ng of input DNA. The tagmentation reaction was quenched, purified, and loaded onto a 0.6% agarose gel as before. DNA fragments between *ca.* 1.7 kb and 6.3 kb were collected and purified by gel extraction as above, eluting twice with 6 μl of 55°C water. To the 12 μl elution was added 12 μl of 2X Q5 master mix previously held at 98°C for 30 seconds and the gap-filling reaction was held at 72°C for 15 minutes before purification and elution, resulting in 60.5 ng of blunt ended DNA fragments. All 60.5 ng of insert DNA (∼0.05 pmol assuming 2 kb average size) were added to a 20 μl NEBuilder HiFi assembly reaction with 35 ng of linear pZE21-ME vector (∼0.025 pmol) and held at 50°C for 15 minutes followed by column purification and elution twice with 6 μl of 55°C water. The purified assembly reaction was transformed into electrocompetent *E. coli*, rescued, amplified, titered for cfu count, and subjected to colony PCR to determine average insert length and library size as above.

### Extraction of plasmid DNA from un-selected libraries

Functional metagenomic library stocks prepared from strains ABC07 and ABC10 (see above) were plated in triplicate on LB+KA50 agar to determine cfu/ml concentrations. Each triplicate library was plated again on LB+KA50 agar plates with volumes calculated to result in ∼1000 colonies plated, resulting in an average of ∼600 colonies per plate following overnight incubation at 37°C. Colonies on each plate were collected by addition of 750 μl of LB to the plate and resuspending colonies using a cell spreader and removal of the media. This process was repeated, resulting in ∼1 ml of turbid bacterial culture which was subsequently used as input for plasmid purification by miniprep kit (New England Biolabs, cat#T1010S). The pooled plasmid library DNA for each library was eluted in 35 μl of elution buffer and quantified, resulting in an average concentration of 58 ng/μl miniprep.

### Functional metagenomic selection for antibiotic resistance

Libraries to be tested were first plated on LB+KA50 agar plates to determine cfu/ml titer following which the volume of frozen stock necessary to provide 10-fold coverage of unique inserts in each library was calculated. This volume, brought up to 100 μl with LB media if necessary, was plated on Mueller-Hinton II cation adjusted agar plates containing 50 μg/ml kanamycin (MH+KA50) and another selective antibiotic depending on the library. Triplicate ABC07/ABC10 libraries prepared by METa assembly or blunt ligation were plated on MH+KA50 supplemented with 1,000 μg/ml (1 mg/ml) penicillin G sodium salt (Fisher Scientific, cat#AAJ6303214). The 162 Gb soil library was plated on MH+KA50 supplemented with 64 μg/ml nourseothricin sulfate (Dot Scientific, cat#DSN51200-1) and the 300 ng goose fecal pellet library was plated on MH+KA50 supplemented with either 8 μg/ml tetracycline (Fisher Scientific, cat#AAJ6171406) or 4 μg/ml colistin sulfate (Fisher Scientific, cat#AAJ6091503). Following overnight selection of plates at 37°C resistant colonies were collected as slurries and plasmids extracted and purified as above. Antibiotic concentrations were chosen based on literature precedent (*i.e.* 1 mg/ml penicillin (47, 48), 8 μg/ml tetracycline (37), 4 μg/ml colistin (35), and 64 μg/ml nourseothricin (52)). Complete growth inhibition of *E. coli* DH10B with empty pZE21-ME vector was confirmed for each antibiotic after plating a similar high density lawn and incubating at 37°C overnight.

### Amplicon sequencing of functional metagenomic libraries

Plasmid minipreps from un-selected and antibiotic selected libraries were used as templates for PCR reactions targeting vector inserts. Seven 100 μl PCR reactions, one for each library, were performed containing 50 μl of Q5 2X master mix, 39 μl water, 5 μl each of 10 μM primers 6463TSC and 6464 TSC, and 1 μl of library miniprep corresponding to between 5.4 ng and 8.6 ng of DNA. Reactions were run in a thermocycler using the following settings: holding at 98°C for 30 seconds followed by 16 cycles of 98°C for 10 seconds and 72°C for 4 minutes followed by holding at 72°C for 5 minutes. Reactions were purified by silica column and each column was eluted twice with 20 μl of 55°C water and quantified. Insert amplicon integrity was verified by running 100 ng of purified DNA from each reaction on a 0.8% agarose gel stained with SYBRSafe.

Samples were shipped overnight to the University of Illinois at Urbana-Champaign Roy J. Carver Biotechnology Center as 30 μl aliquots containing 500 ng of each reaction. Samples were used as input for library preparation and sequencing on the PacBio Sequel II platform at the center as follows. Amplicons were ligated to barcoded adaptors using the Barcoded Overhang Adaptors Kit (Pacific Biosciences, CA). The barcoded amplicons were normalized to the estimated number of unique inserts, based off of colony counts, and pooled. The pooled amplicons were used as input for a SMRTBell Express Template Prep kit 2.0 (Pacific Biosciences) to prepare the sequencing library. The library was quantitated by Qubit fluorometer, and DNA fragment size and quality were confirmed on a Fragment Analyzer (Agilent, CA). The library was sequenced on a SMRTcell 8M on a PacBio Sequel II with 20 hs movie time. Circular consensus analysis was performed on the resulting BAM file using SMRTLink V8.0 using the following parameters: ccs -min-length 500 --max-length 12000 --min-passes 3 --min-rq 0.99.

### Demultiplexing was performed with lima (Pacific Biosciences) using default parameters. Analysis of functional metagenomic library sequencing

Read coverage and evenness of the ABC07/ABC10 libraries were determined for each genome individually. Long reads were mapped to either ABC07 (ASM217990v1) or ABC10 (ASM217996v1) assemblies using minimap2 v2.17-p94 with default settings, converting resulting SAM files to BAM files with samtools v1.9 (53) using default settings, and calculating the average read coverage over a 1,000 bp window using pileup.sh from the BBmap v38.86 suite. For visualization of coverage, a rolling average was produced using the rollmean() function from the Zoo R package v1.8.8 (54).

To identify potential β-lactamase genes in the ABC07 (ASM217990v1) and ABC10 (ASM217996v1) genomes the amino acid fasta files were downloaded and individually submitted to the Comprehensive Antibiotic Resistance Database (CARD) Resistance Gene Identifier (RGI) webserver (https://card.mcmaster.ca/analyze/rgi) (55) for prediction of all resistance genes using parameters ‘Protein sequence’, ‘Perfect, Strict and Loose’, ‘Include nudge’, and ‘High quality/coverage’. The resulting dataframe was filtered for antibiotic resistance genes belonging to the β-lactamase AMR Gene Family that had ‘antibiotic inactivation’ as its resistance mechanism to generate a list of ABC07 or ABC10-specific β-lactamase amino acid sequences. For each functional metagenomic approach and selection method, the following was performed on all reads that mapped to either ABC07 or ABC10. Open reading frames (ORFs) were identified using Prokka v1.14.6 (56) with parameters –norrna, --notrna, --noanno, and --fast. Resulting ORFs for each combination were clustered with mmseqs2 linclust (57) with parameters --min-seq-id 0.95 and -c 0.95. Predicted β-lactamases were searched against the set of representative ORFs using an e-value cutoff of 10^-6^. Afterwards, a single best hit determined by bitscore, identity, and coverage was kept. A hit was considered a true β-lactamase if it had ≥95% coverage and identity.

Functional annotation of the soil and goose fecal microbiome libraries was carried out by ORF searching and clustering all metagenomic long reads using Prokka and mmseqs2 linclust as previously described. Representative ORFs were submitted to the CARD RGI portal to search for ARGs using the previously stated parameters. Non-resistance functional annotations were obtained by submitting representative ORFs to eggNOG-mapper v2 (http://eggnog-mapper.embl.de/) (58). Antibiotic resistance genes and functional annotations were then mapped back to all identified ORFs using a custom script. Counts of antibiotic resistance genes, antibiotic resistance gene ontology, superfamilies, and drug class were extracted from the CARD RGI output.

Mobile genetic elements syntenic to antibiotic resistance genes were identified based on keyword searches of the eggNOG ‘annotation’ column using the search terms “transposase”, “conjugative”, “phage”,’ “integrase”, “replication”, and “recombinase”. ORFs that matched these search terms were further verified using UniProt (https://www.uniprot.org/) (59). Plots were made using R package ggplot v3.3.2 (60) and gene region visualizations were created using R package gggenes v0.4.0.

Streptothricin acetyltransferase gene family phylogenetic analysis was performed by extracting the five amino acid sequences for streptothricin resistance enzymes from the CARD database and using these as input to run NCBI blastp (61) against the NR database (run November, 2020). Sequence hits were filtered for ≥25% amino acid identity and ≥70% alignment length to match CARD workflow. Replicates were removed from the combined sequences followed by clustering using CD-Hit (62) at a 90% identity threshold. The amino acid sequences of three representative putative streptothricin acetyltransferase enzymes from the nourseothricin selection were added to the sequence file and all were aligned using mafft v7.471 (63) with parameters –thread 8 –localpair –maxiterate 1000. An approximate maximum likelihood phylogenetic tree was calculated using FastTree v2.1.11 (64) with parameter-wag. The resulting phylogenetic tree was visualized with ggtree (65).

### Literature search for comparable library statistics

A literature search was carried out on the National Library of Medicine using PubMed using the search terms “functional metagenomics” OR “metagenomic libraries” on September 30^th^, 2020. The results were sorted by publication date and the 125 most recent publications were manually searched for functional metagenomic library preparation details including insert size (small or large), input DNA mass, and total library size. An additional three publications known to contain these details were appended to the 125 publications to give a total of 128 publications searched. Off-topic publications, publications that do not describe the preparation of new libraries, and publications with insufficient methods details were removed leaving six suitable publications: Campbell *et al.* (2020), Gasparrini *et al.* (2019), Kintses *et al.* (2019), Marathe *et al.* (2018), Gibson *et al.* (2016), and Moore *et al.* (2013) (21, 28, 35, 66–68). Library efficiency was determined by normalization of reported library size to reported or best estimate input DNA mass.

## RESULTS

### Strategies to improve functional metagenomic library preparation efficiency

Our first strategy to increase functional metagenomic library preparation efficiency was to replace sonication-based fragmentation with transposase-based tagmentation (**Figure 2A**). In theory, this would allow for lower input DNA mass and remove expensive capital equipment from the blunt ligation cloning protocol (**Figure 2B**).

Our second strategy to increase efficiency was to replace blunt ligation cloning with homology-based seamless assembly cloning, taking advantage of the fact that tagmentation-produced fragments would all have known DNA sequence on their ends. One commercial assembly protocol that we hypothesized to be compatible with tagmented DNA fragments was In-Fusion cloning from Takara (69). This method is optimized for overlap regions with lengths between 12 bp and 21 bp and is therefore compatible with homology consisting of 19 bp mosaic end sequences. In-Fusion uses 3’ exonuclease activity to produce single stranded DNA 5’ overhangs, allowing treated inserts and vector to hybridize. In-fusion assembly does not result in a covalently closed circular DNA product but instead retains gaps and nicks that are resolved post-transformation in the recipient cell (**Figure 2C**).

A second assembly option that we hypothesized to be compatible was NEBuilder HiFi DNA assembly from New England Biolabs, which functions similarly to Gibson assembly (44). This method requires overlap regions with melting temperatures greater than 50°C which is compatible with the 19 bp mosaic end sequence (an estimated melting temperature of 52°C by 2AT+4GC rule). NEBuilder HiFi and Gibson assembly use 5’ exonucleases to produce 3’ overhangs. Because tagmentation results in covalent addition of mosaic end oligos to only the 5’ ends of DNA fragments, 5’ exonuclease activity would effectively erase the mosaic end sequence from the inserts. Nextera tagmentation protocols overcome this obstacle by including a brief DNA polymerase gap filling reaction that would be applicable to our protocol as well. The resulting DNA would be mixed with the NEBuilder HiFi assembly master mix, resulting in 3’ overhangs able to hybridize to complementary sequences on linearized vector. Following hybridization, NEBuilder HiFi assembly master mix polymerase and ligase fill gaps and ligate nicks, respectively, to produce a covalently sealed construct for transformation (**Figure 2D**).

### Expression and purification of transposase enzyme and 1 kb to 10 kb DNA tagmentation

We began our experiments with the goal of increasing DNA efficiency during metagenomic library preparation by adapting the tagmentation work flow used in shotgun sequencing library protocols (38, 39). This was aided by the availability of non-commercial protocols for the preparation of the transposase reagent (41, 42). We modified these published protocols for expression and purification of Tn5 transposase enzyme, most notably in our use of auto-induction media (45) and other expression conditions. Our expression system responded well to auto-induction (**Supplemental figure 1A**) with one liter of culture yielding 8 g of wet cell mass and we obtained approximately 10.5 mg of well purified enzyme (**Supplemental figure 1B**) in good agreement with reported yields of 15 mg per liter of culture (41). We verified the activity of our transposase by observing its ability to convert metagenomic DNA into low molecular weight fragments <500 bp in size (**Supplemental figure 1C**).

Next, we investigated if transposase could be used to prepare metagenomic or genomic DNA in fragments capable of containing bacterial open reading frames (ORFs, roughly 1 kb to 5 kb in length). We used high molecular weight genomic DNA (measured at ∼70 kb) isolated from two penicillin-catabolizing bacteria, ABC07 (*Pseudomonas* sp. PE-S1G-1) and ABC10 (*Pandoraea* sp. PE-S2T-3) (46–48) as input for these test reactions. While tagmentation is usually used to create ∼200 bp DNA fragments for sequencing on the Illumina platform we were able to alter this by adjusting the ratio of transposase to input DNA (similarly, others have successfully used another enzyme used to prepare shotgun sequencing libraries, fragmentase, to make functional metagenomic libraries (67)). Using 0.5 ng of transposase per ng of target DNA yielded a fragmentation pattern centered around 2.5 kb (**Figure 3**).

**Figure 3.**
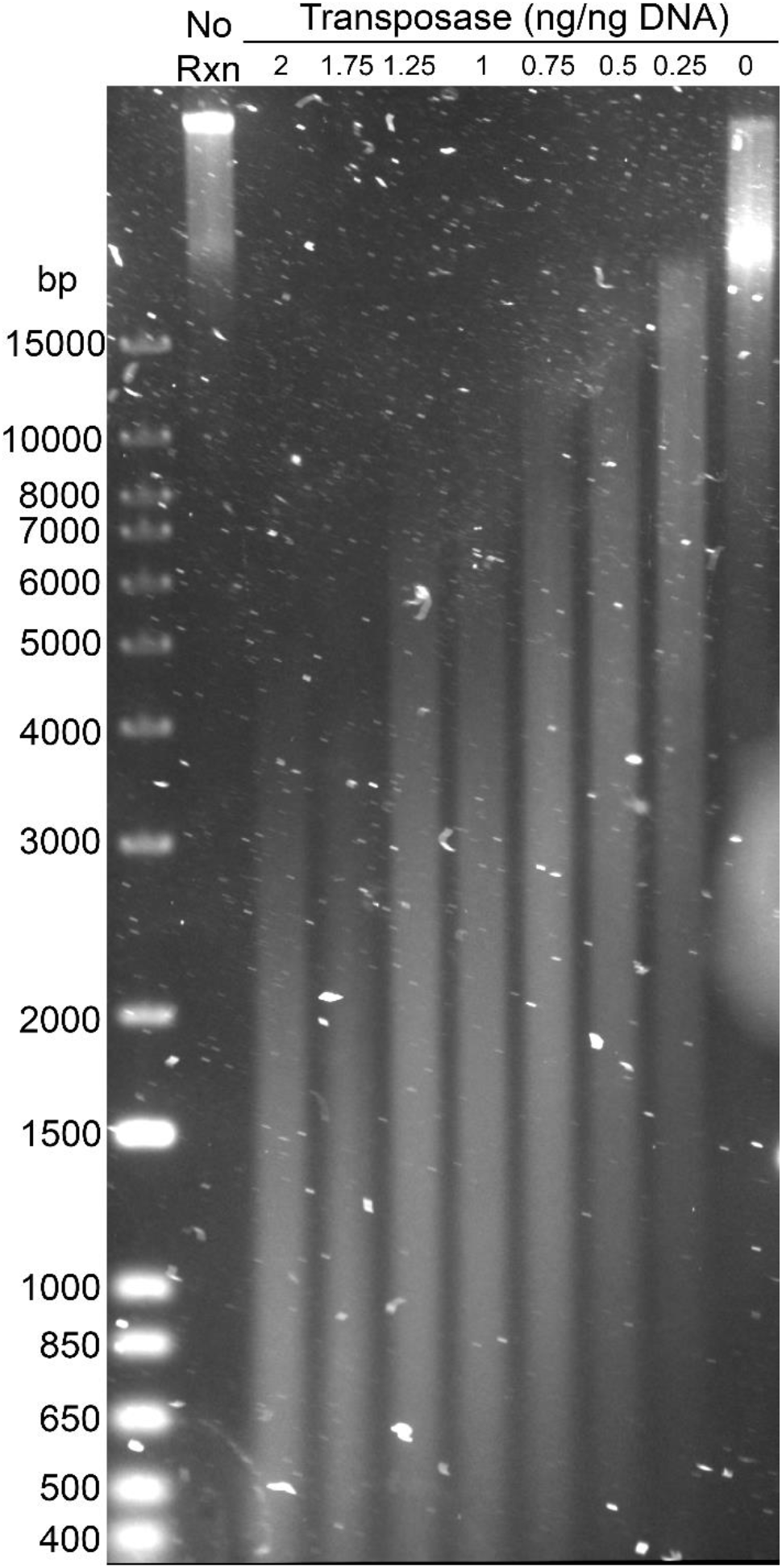
Transposase to target DNA ratios for 1 kb to 10 kb insert fragmentation. High molecular weight genomic DNA (no rxn = no reaction) was incubated with transposase at concentrations from 0 ng/ng of DNA up to 2 ng/ng DNA and analyzed by pulsed field agarose gel electrophoresis.

### METa assembly using In-fusion or NEBuilder HiFi assembly compared to blunt ligation

We hypothesized that inserts prepared by tagmentation could be compatible with both blunt ligation and assembly-based methods of functional metagenomic library preparation. To test this hypothesis, we extracted high molecular weight DNA from a soil sample and fragmented it by tagmentation, resulting in fragments between approximately 1 kb and 8 kb in length (**Supplemental figure 2**). We purified the fragmented DNA by excision from an agarose gel and used the resulting pool of mosaic end 5’ tagged fragments as input for three library preparation methods, described above (**Figure 2**), in triplicate. DNA fragment preparation began with a single aliquot of 4.5 μg of metagenomic DNA which dropped to 3.175 μg following tagmentation and PCR purification (∼30% loss), and dropping to 2.145 μg of DNA following size selection and gel purification (∼30% step loss, ∼50% loss overall). After cloning inserts into vector (by enzymatic assembly or ligation), we transformed the entirety of each purified reaction by electroporation into homemade competent *E. coli* cells. We plated dilutions of the recovered cells to determine titer. The following day, we used colony PCR to find the average insert size and proportion of colonies with an insert (as opposed to empty vectors) (**Supplemental figure 3**).

METa assembly using In-Fusion premix (**Figure 2C**) resulted in recovery of only a single colony on a 100-fold dilution plate and we did not pursue this strategy any further. We instead focused on comparing libraries created *via* blunt ligation to libraries created by METa assembly with NEBuilder HiFi. Blunt ligation cloning (**Figure 2B**) resulted in, on average, dozens of colonies on agar plates spread with 100-fold diluted transformation culture corresponding to 1.6×10^4^ cfu, 4.6×10^4^ cfu, and 5.2×10^4^ cfu for each replicate. In comparison, NEBuilder HiFi mediated METa assembly (**Figure 2D**) agar plates had uncountable colonies at 100-fold dilutions and ∼100 colonies on petri dishes plated with 10,000-fold dilutions, corresponding to 1.3×10^6^ cfu, 9.8×10^5^ cfu, and 4.9×10^5^ cfu for each replicate. When we normalized these plate counts to how much insert DNA was used during triplicate cloning reactions we found that the NEBuilder HiFi reactions resulted in ∼17.5-fold higher cfu/ng assemblies and an associated *p-*value of 0.0205 (**Figure 4A**). Average insert length did not appear to differ between the two methods, with the average NEBuilder HiFi assembly insert being 0.94 times as large as the average blunt ligation insert (**Figure 4B**). NEBuilder HiFi assembly reactions resulted in significantly larger libraries (∼30-fold larger, *p* = 0.0235) (**Figure 4C**), in part because of their higher titer (**Figure 4A**), but also because all of the colonies assayed contained inserts whereas approximately half of the blunt ligation colonies carried empty vectors or insert DNA <200 bp in length despite both methods using the same linearized vector DNA (**Supplemental figure 3**).

**Figure 4.**
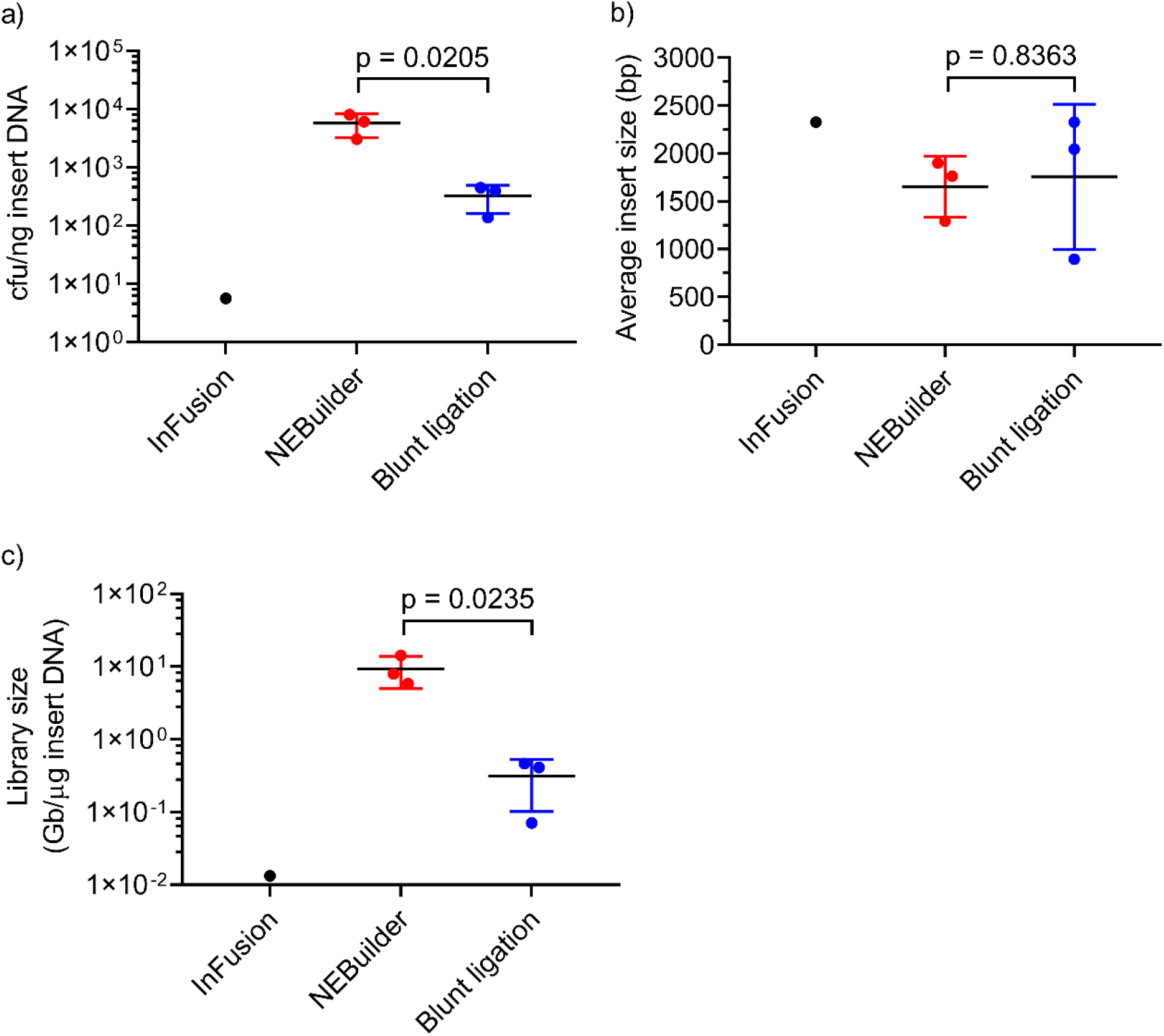
Comparing metagenomic libraries prepared by assembly or blunt ligation. Functional metagenomic libraries were created using METa assembly *via* In-Fusion assembly or NEBuilder HiFi assembly and compared to a library constructed using blunt ligation. All libraries used the same input DNA and were each prepared in triplicate. Error bars represent standard deviation of n=3 experiments. Comparisons between blunt ligation and NEBuilder HiFi METa assembly were made using unpaired two tailed t tests. Only a single In-Fusion colony was isolated and therefore we did not perform statistical tests on that method. a) Post-transformation culture titers normalized to the quantity of insert DNA used in the assembly or cloning reaction itself. b) Average insert size for plasmids containing an insert. Colony PCR was performed on the single In-Fusion colony, and 9 colonies were analyzed and averaged for each replicate NEBuilder HiFi assembly or blunt ligation reaction to give three data points per method. c) Final total library size or each replicate measured in gigabase pairs (Gb) of captured metagenomic DNA normalized to the amount of insert (μg) used during cloning or assembly.

### NEBuilder HiFi METa assembly compared to blunt ligation

Because of the failure of our In-Fusion mediated METa assembly reactions, we focused on NEBuilder HiFi mediated METa assembly reactions which will be referred to simply as METa assembly from this point forward. The apparent much higher efficiency of METa assembly compared to traditional blunt ligation in constructing metagenomic libraries (**Figure 4**) led us to seek replication of our findings. Our approach was broadly similar to that detailed above, with two changes. First, instead of using soil metagenomic DNA as input to tagmentation, we combined and tagmented (**Supplemental figure 4**) genomic DNA of two bacterial strains of interest to us: *Pseudomonas* sp. Strain PE-S1G-1 (ABC07) and *Pandoraea* sp. Strain PE-S2T-3 (ABC10). These previously sequenced strains (46) have been shown to be capable of using the antibiotic penicillin as their sole carbon source (48) *via* a pathway likely initiated by a β-lactamase enzyme (47). Creation of a functional mixed genomic library from these strains would allow us to characterize penicillin resistance genes in a sequence naïve manner and test previous annotations and findings. Second, instead of transforming METa assembled or blunt ligation cloned libraries into homemade electrocompetent cells we purchased electrocompetent *E. coli* cells with much higher transformation efficiency. These are routinely used in functional metagenomic library preparation and result in larger libraries (37).

We found that METa assembly again resulted in significantly higher titers of transformed cells per ng of insert used during library preparation (**Figure 5A**, ∼276-fold greater than the blunt ligation libraries, *p* = 0.038) and significantly larger libraries (**Figure 5C**, ∼296-fold greater than the blunt ligation libraries, *p* = 0.006). Colonies containing empty vectors occurred much more frequently following blunt ligation reactions (3/7, 4/8, and 5/9 by colony PCR) compared to colonies from METa assembly reactions (0/7, 0/7, and 0/8 by colony PCR) (**Supplemental figure 5**). Average insert size (**Figure 5B**) did not appear to differ significantly between methods.

**Figure 5.**
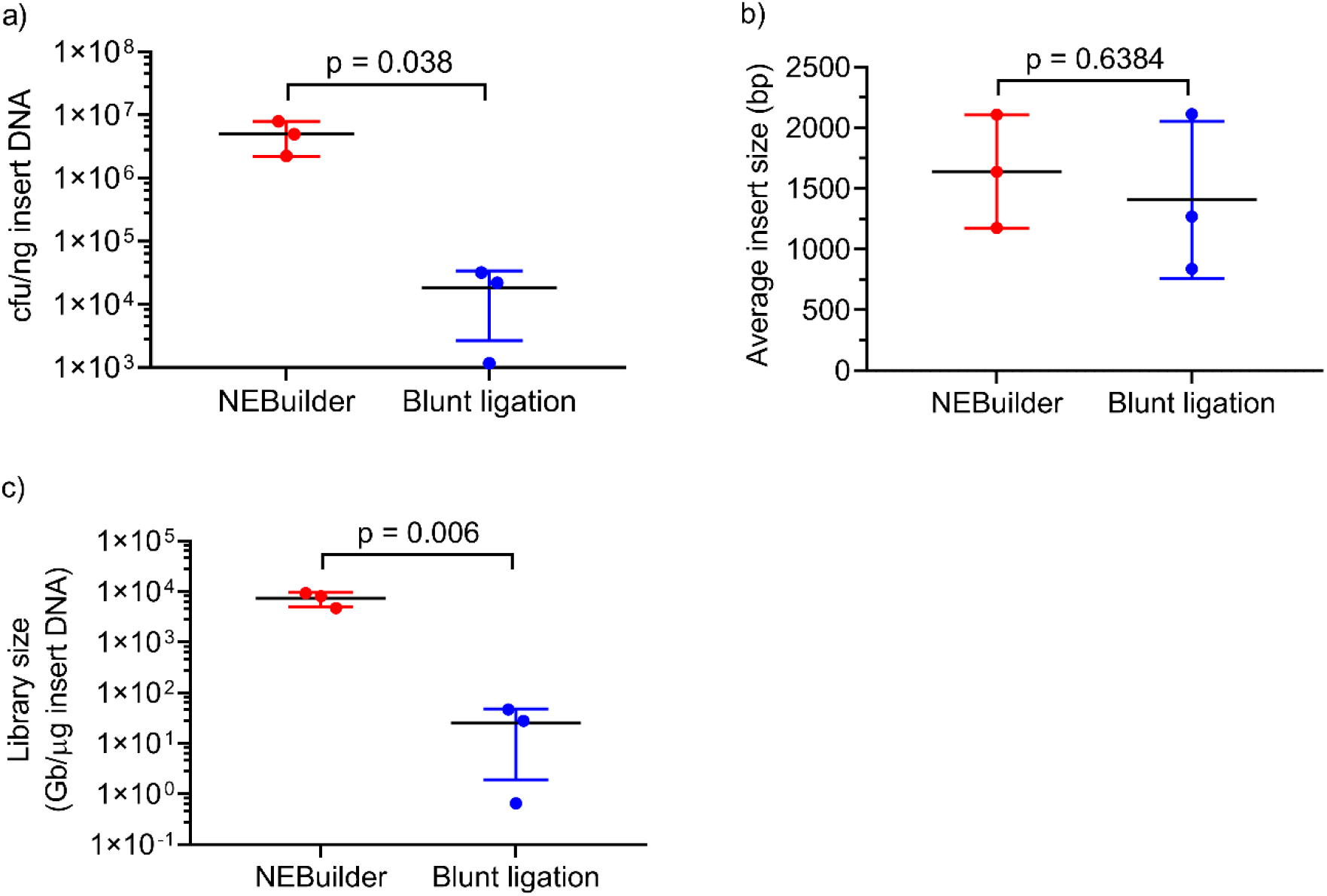
Libraries created by NEBuilder HiFi mediated METa assembly are larger than those created by blunt ligation. Both sets of triplicate metagenomic library assembly/cloning reactions used the same input DNA and were compared using unpaired two tailed t tests. Error bars represent standard deviation of n=3 experiments. a) Culture titers of recovered cells post-transformation normalized to insert DNA mass used in assembly or cloning. b) Vector average insert size determined by colony PCR, excluding colonies containing empty vector constructs. 13 colonies were assayed and insert lengths were averaged for each replicate (n=3) for each construction method. c) Final library size given in Gb/μg of insert DNA.

After establishing that assembly cloning results in much larger libraries than blunt ligation cloning we next asked if genomic coverage was similar across methods. We sequenced between 355 and 831 random colonies from each of the six libraries. Each library was plated to give approximately 1,000 colonies based on prior titers resulting in an average of approximately 600 colonies collected (*ca.* 1562 total colonies from blunt ligation plates, 2008 total colonies from assembly plates). We extracted plasmids from the collected colonies and pooled each set of triplicate libraries into a single pool. These two pools were used as templates in a limited PCR reaction to amplify inserts (**Supplemental figure 12**) which were submitted for long read sequencing on the PacBio Sequel II platform. The resulting reads were mapped back onto published ABC07 and ABC10 genomes (46) and the nucleotide coverage of each library for each genome was calculated. We found qualitatively good agreement between assembly and blunt ligation library coverage of both ABC10 (**Figure 6A**) and ABC07 (**Supplemental figure 6A**).

**Figure 6.**
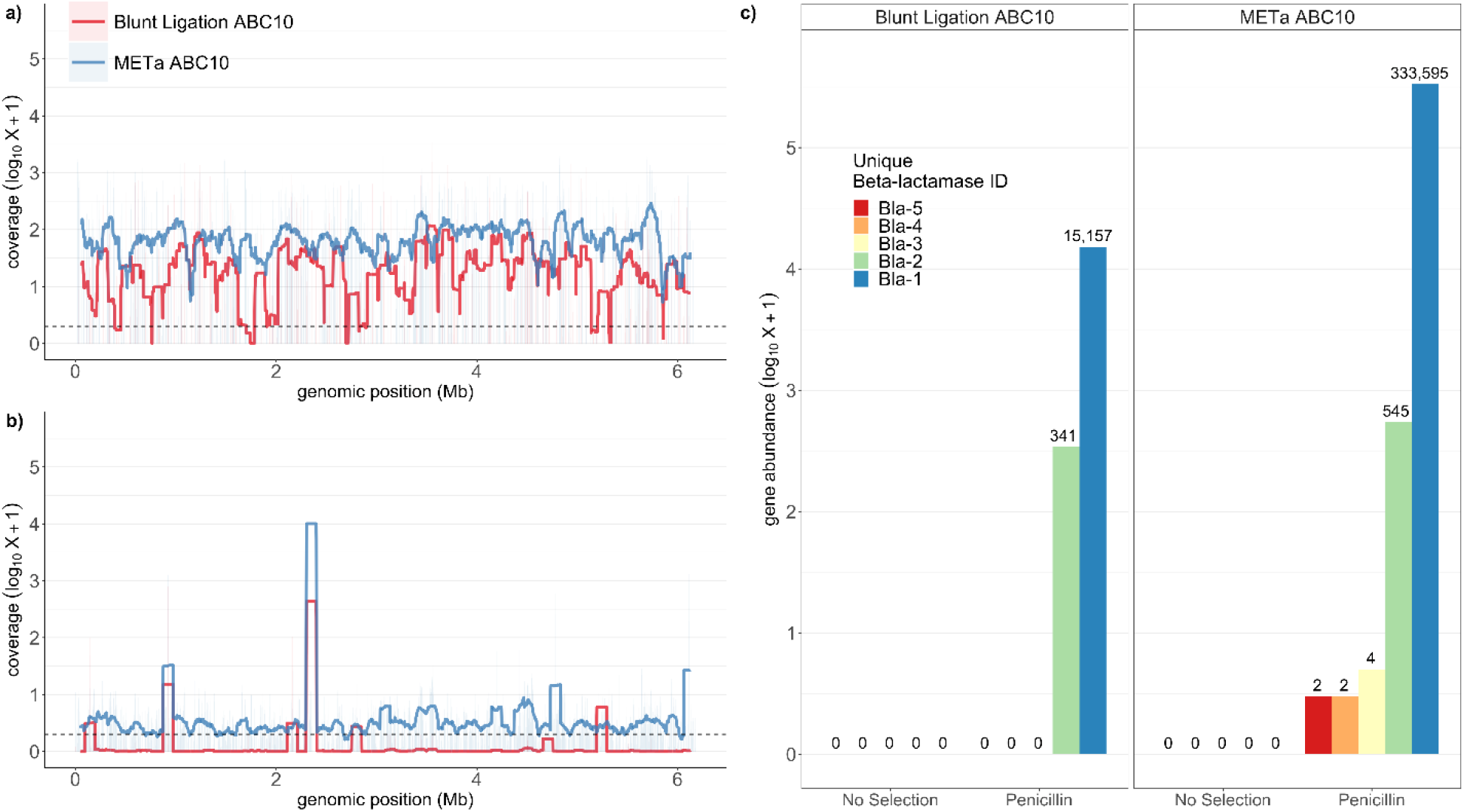
Assembly and blunt ligation library coverage of ABC010 genome with and without penicillin selection. a) Nucleotide depth of coverage for ABC10 genome by functional metagenomic library prepared by assembly (blue) or blunt ligation (red). Coverage is smoothed to a 1 kb resolution. b) Same as in a) but sequenced libraries were first subjected to selection on agar plates containing 1 mg/ml penicillin. c) Gene abundance in post-penicillin selection reads for each of five predicted ABC10 β-lactamase genes that had read number >0. From Bla-1 to Bla-5 respective NCBI accession numbers are WP_087722475.1, WP_087721859.1, WP_140413467.1, WP_087721948.1, and WP_087721885.1.

In order to verify the functional aspect of our functional metagenomic libraries we next performed triplicate selections for growth in the presence of 1 mg/ml penicillin. This concentration, about 10-fold higher than what is normally used in microbiology, was chosen based on high level penicillin resistance of strains ABC07 and ABC10. Stocks were plated with the goal of plating enough cells to reach 10-fold coverage of each library, resulting in denser plating for the much larger METa assembly triplicate libraries. As before, colonies were collected for PCR amplification of inserts (**Supplemental figure 12**) and sequenced. Sequencing for both libraries were dominated by reads mapping to one region (ABC07, **Supplemental figure 6B**) or two regions (ABC10, **Figure 6B**) of the donor organism genomes. In each case these regions corresponded to predicted β-lactamase genes (**Figure 6C, supplemental figure 6C**).

### Preparation of a large soil metagenomic library by METa assembly

We next supplemented our method comparisons (**Figures 4 and 5**) by preparing a functional metagenomic library by METa assembly for direct comparison to published libraries prepared by blunt ligation. A common DNA input quantity for preparation of a single functional metagenomic library by the sonication/blunt ligation workflow is ∼5 μg (21, 28, 35, 37, 66, 70). In order to compare METa assembly to the broader literature, we performed a tagmentation reaction on 5 μg of soil metagenomic DNA. Following size selection and polymerase gap filling we retained 1.122 μg of mosaic end containing inserts ready for assembly. Because this quantity of DNA falls well outside the recommended capacity of NEBuilder HiFi reactions we proceeded to first assemble 16.6% (175 ng) of the total in a trial reaction. Transformation of the purified assembly reaction resulted in a 162 Gb library (7.8×10^7^ unique clones, average insert size 2.077 kb ± 708.7 bp standard deviation, **Supplemental figure 7**). After scaling up assembly and transformation, the remaining inserts were used to prepare a 529.5 Gb library (2.85×10^8^ unique clones, average insert size of 1.858 ± 555.1 bp). Combined these libraries form a 691.5 Gb library from 5 μg of input DNA.

To test the utility of this method, we performed a functional metagenomic selection on the 162 Gb library using the natural product antibiotic nourseothricin. Nourseothricin (also known as cloNAT) is a member of the streptothricin class of aminoglycoside antibiotics first described by Waksman and Woodruff in 1942 (71). Insert amplification (**Supplemental figure 12**) and sequencing resulted in identification of acetylation to be the dominant mode of resistance in our library (**Supplemental figure 8A**). Among these we identified multiple apparent homologs to known streptothricin acetyltransferases, with some inserts encoding multiple syntenic resistance determinants (**Figure 7A**). Phylogenetic analysis of nourseothricin acetyltransferase proteins places our representative soil-derived resistance genes within this family, with one representative appearing to represent a novel cluster related to the StaT enzyme (26.22% identity, 96.83% coverage) and two representatives clustering near SatA and Sat-4 enzymes (50.28%/47.96% identity and 102.17%/95.11% coverage) (**Figure 7B**). This clustering pattern is in agreement with the amino acid sequence percent identities between the soil-derived acetyltransferases and CARD validated streptothricin resistance enzymes.

**Figure 7.**
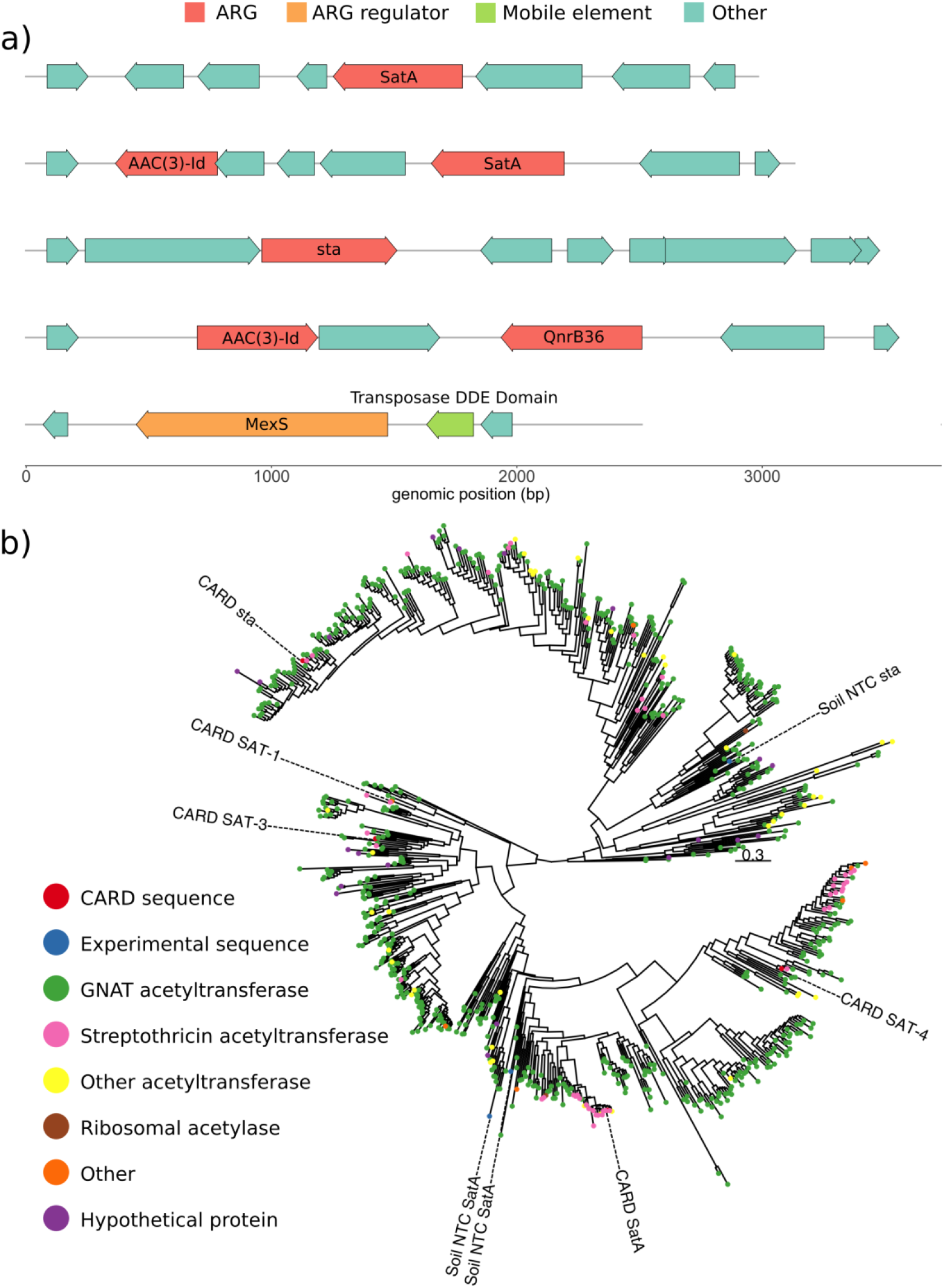
Nourseothricin resistance conferring inserts from soil microbiome library selections. a) Genomic context of representative nourseothricin resistance genes including syntenic mobilization or regulatory elements and other antibiotic resistance genes. b) Phylogenetic tree of five CARD streptothricin acetyltransferase enzymes (red circles, [CARD]), three soil metagenome nourseothricin resistance genes (blue circles, [Soil NTC]), and related enzymes.

### METa assembly of functional metagenomic libraries using limited input DNA

Our experiments comparing METa assembly library creation against library creation using blunt ligation suggest that METa assembly, with its potential for low input DNA mass and higher efficiency, may allow for the creation of functional metagenomic libraries using samples with low DNA mass or quality. We first tested this possibility by creating a mock sub-optimal metagenomic DNA sample consisting of a 10 kb λ phage DNA amplicon (in place of high-quality DNA >48 kb suggested for Covaris sonicator). We used 200 ng of the DNA as input into a tagmentation reaction (in place of the suggested 2 μg to 20 μg suggested for the sonicator). Following clean-up, size selection, and polymerase repair we obtained 59.4 ng of 1 kb to 5 kb DNA fragments, all of which were used as input for an assembly reaction. This resulted in a ∼13.5 Gb library with average insert size of ∼1.02 kb ± 340 bp standard deviation (**Supplemental figure 9**) and a library efficiency of 67.5 Gb/μg of input DNA, suggesting that METa assembly can still work well with low input and lower quality (*i.e.* not high molecular weight) DNA.

Because it is possible that amplicon DNA behaves differently from metagenomic DNA, we next performed a low input METa assembly library preparation using the same soil metagenomic DNA as in the 5 μg library. This time we started with a 250 ng DNA input tagmentation reaction and after clean-up, size selection, and polymerase repair we performed an assembly reaction using all 35.2 ng that passed through the processing steps. Electroporation resulted in 3.8×10^6^ colonies with an average insert size of 2.029 kb ± 284 bp standard deviation (**Supplemental figure 10**). We calculated the library size to be 7.71 Gb and the library efficiency to be 30.84 Gb/μg, suggesting that METa assembly retains high efficiency with low input metagenomic DNA. Because we had prepared the previous soil library we did not test this library any further.

Finally, we prepared one additional low input DNA mass library by METa assembly. We chose to use metagenomic DNA extracted from a Canada goose fecal pellet in order to include DNA sourced from a non-soil microbiome and due to its ubiquitous availability near our research space (**Supplemental figure 11A**). We used 300 ng of input DNA for tagmentation and, following the usual sample processing steps, performed assembly into vector using 60.5 ng of size-selected and repaired inserts. The resulting library was estimated to contain 1.18×10^7^ unique clones with colony PCR indicating 95.2% of clones containing an insert, with an average insert length of 2.39 kb ± 651.5 bp standard deviation (**Supplemental figure 11B**). Together, this suggests a total library size of 26.86 Gb and a library efficiency of 89.67 Gb/μg.

We selected the goose fecal microbiome library against a widely used soil natural product antibiotic, tetracycline, and an antibiotic of last resort, colistin. Metagenomic inserts were amplified from functionally selected plasmids by PCR (**Supplemental figure 12**) and sequenced. The dominant mechanism for tetracycline resistance in the goose gut microbiome appears to be drug efflux (**Supplemental figure 8C**). Many of the genes encoding efflux pumps are syntenic to known regulatory elements (*e.g. tetR*) and/or potential mobilization elements (*e.g.* transposase or phage integrase genes) (**Figure 8A**). The dominant mechanisms for colistin resistance in our library consist of modification of lipid A and antibiotic efflux (**Supplemental figure 8B**). Among the predicted lipid A modifying enzymes we identified homologs to the emerging MCR family of mobilized colistin resistance enzymes, including an MCR-5.2 homolog (36.14% identity, 109.89% coverage to CARD representative) (**Figure 8B**).

**Figure 8.**
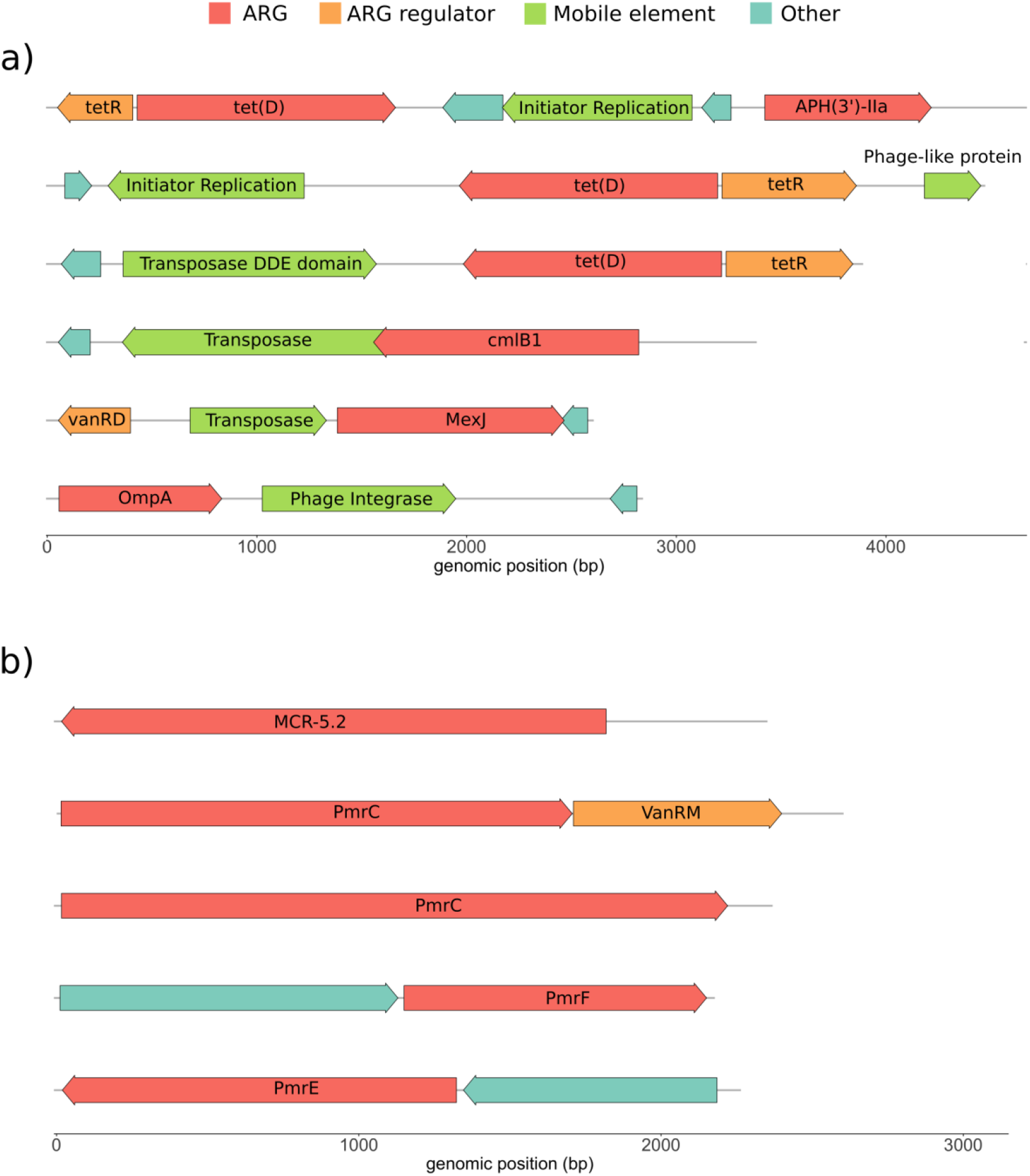
Antibiotic resistance conferring inserts from goose gut microbiome library selections. Genomic context of representative resistance genes including syntenic mobilization or regulatory elements and other antibiotic resistance genes following goose microbiome library selection on a) tetracycline or b) colistin.

## DISCUSSION

Functional metagenomic libraries have been valuable tools in chemical biology and have been instrumental in the discovery of novel enzymes involved in antibiotic biosynthesis and resistance, pharmaceutical-microbiome interactions, bioremediation of pollutants, and many other activities of value to medicine and industry (15–17). However, the methods used in the preparation of small insert functional metagenomic libraries (**Figure 1**), sonication fragmentation and blunt ligation cloning, currently limit library size, efficiency, and adoption of this method. DNA fragmentation by sonication is widely used because it shows low bias and it is possible to control the resulting fragment size distribution. However, sonication instruments represent a significant capital expense and require a high mass of high quality input DNA (*e.g.* from 2 µg to 20 µg of >48 kb length for a Covaris sonicator). This has limited the microbiomes that can be explored by functional metagenomic libraries to those with a large amount of available metagenomic DNA. Blunt ligation, in turn, uses input DNA inefficiently due to the low probabilities involved in the random collisions that bring blunt DNA ends and ligase together to react. Significant input DNA is also wasted by requiring a five-fold molar excess of input DNA to vector DNA to avoid vector self-ligation. Due to this low efficiency, upwards of 20 μg of input DNA may be recommended for library preparation (37). This is illustrated by a recent publication (21) where the authors prepared 22 functional metagenomic libraries with an average size of 18 Gb using approximately 5 µg of metagenomic DNA for each library. As 1 μg of input DNA theoretically represents just under 1 million Gb of DNA, it is self-evident that only a small fraction of the input metagenomic DNA is successfully captured. Two additional considerations also arise from the use of blunt end ligation in current forms of functional metagenomic library preparation. Blunt ligation reactions often require overnight incubation, a significant loss of time, especially for protocols that propose using functional metagenomics as part of a diagnostics pipeline (25). The use of blunt ends also results in library vulnerability to contamination by foreign DNA. Custom agarose gel ladders lacking bacterial DNA and bleaching and washing of electrophoresis apparatus are required to combat this risk (37).

We were inspired by the use of transposases in sequencing studies us to see if functional metagenomic library preparation could similarly benefit. We found that transposase-mediated fragmentation can be controlled to yield DNA fragments roughly the size of bacterial ORFs (**Figure 3**) with the following empiric conditions for tagmentation reactions being compatible with our enzyme preparation: 10 mM TAPS buffer pH 8.5, 5 mM MgCl_2_, 10% w/v dimethylformamide, input DNA at 10 ng/µl total reaction volume, and assembled transposase at 0.5 ng enzyme per ng of input DNA (*i.e.* 5 ng of assembled transposase per μl). The resulting inserts are compatible with classic blunt ligation based cloning (**Figure 2B**) to produce functional metagenomic libraries (**Figures 4 and 5**) but without any evidence of increased efficiency. Tagmentation of metagenomic DNA was successful across several DNA sources including purified bacterial genomic DNA, PCR amplicons, soil metagenomic DNA, and fecal metagenomic DNA.

We also realized that installation of mosaic end tags on the 5’ ends of fragmented DNA could allow us to replace blunt ligation cloning with higher efficiency homology based assembly methods (**Figures 2C and 2D**). Direct comparisons of functional metagenomic library preparation by assembly cloning demonstrated that METa assembly synergizes dramatically with tagmented DNA fragments and is significantly more efficient (**Figures 4 and 5**) than library preparation *via* blunt ligation. While variation in read counts and the number of input colonies confounded a statistical interpretation, qualitatively it appears that libraries prepared by METa assembly provide equal, if not greater, coverage of input DNA compared to libraries prepared by blunt ligation (**Figure 6A, supplemental figure 6A**).

In order to compare METa assembly against functional metagenomic library preparation by current experts in the field we carried out a literature search of recent articles to find appropriate comparators (**Supplemental figure 13**). Literature protocols with sufficient methods detail (21, 28, 35, 66–68) showed substantial gains in efficiency are made by using METa assembly to prepare functional metagenomic libraries. Literature protocols used initial DNA input masses of between 10 μg and 500 μg to prepare functional metagenomic libraries sized between 10 Gb and 400 Gb. Starting with 5 μg of metagenomic DNA, we used METa assembly to make libraries totaling 712 Gb in size, approximately the same size as the six literature examples combined using a fraction of their total input DNA (**Figure 9A**). Similarly, libraries prepared by METa assembly using only 200 ng to 300 ng of input DNA, more than 10-fold less than the normative 5 μg input for a single library, resulted in libraries of comparable size. When library size is normalized to input metagenomic DNA mass, functional metagenomic libraries prepared by METa assembly significantly outstrip libraries prepared by blunt ligation (**Figure 9B**).

**Figure 9.**
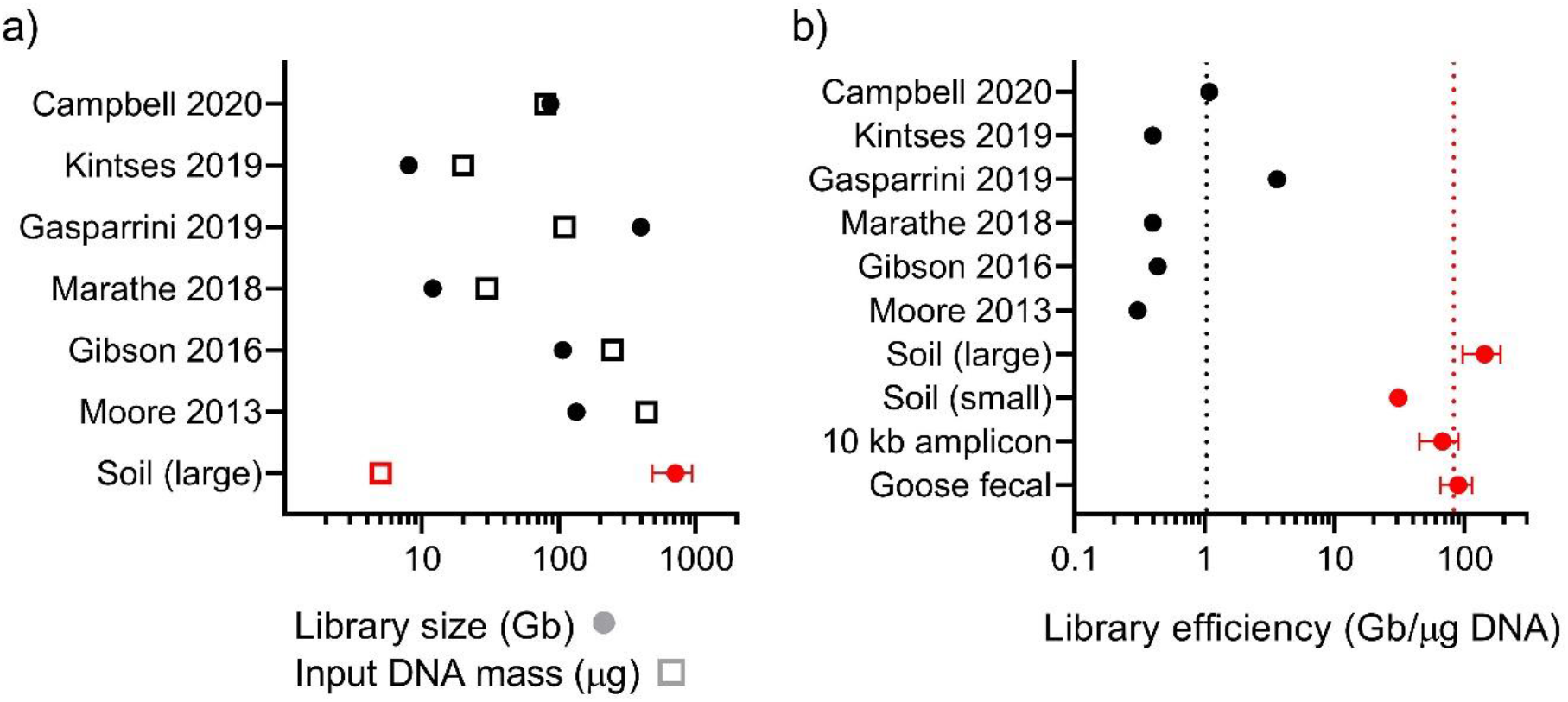
Comparison of METa assembly libraries to literature examples. a) Input DNA mass and functional metagenomic library sizes for six publications (black) compared to the large (5 μg) soil METa assembly library (red). Error bars (not present for literature libraries) calculated based on insert size standard deviation as determined by colony PCR (n=48 colonies). Filled shapes correspond to library sizes and empty shapes correspond to input DNA mass. b) Library efficiency (library size in Gb normalized to input DNA mass in μg) calculated for literature examples (black) or METa assembly libraries (red). Vertical dotted lines correspond to literature (black) or METa assembly (red) average efficiency (n=6 for literature examples, n=4 for METa assembly examples). Standard deviation error bars for METa assembly libraries calculated based on colony PCR as before (from top to bottom colonies tested n=48, 28, 14, or 21).

We next verified the utility of our libraries as chemical biology tools for bioprospecting by subjecting them to selection on three antibiotics. For our soil library we chose to select with the streptothricin antibiotic nourseothricin. Surveys of antibiotic-producing bacteria have determined that the streptothricins are among the most commonly produced antibiotics in soil ecosystems, with between 10% and ∼42% of soil actinomycetes potentially being producers (72, 73). Surprisingly for such a widespread antibiotic there are a limited number of described resistance determinants (*sat2-4*, *sttH*, and *nat1*), especially compared with other aminoglycoside antibiotics (74–76). Our selection and subsequent analysis suggest that the streptothricin acetyltransferase family is larger and more diverse than currently thought and likely contains unknown major branches. These branches likely incorporate many enzymes which at the moment are solely annotated as Gcn5-related N-acetyltransferases (GNATs) (**Figure 7B**). For selection of our goose gut microbiome library we chose to select on two additional classes of antibiotics (in contrast to aminoglycosides and β-lactams used above): tetracycline due to its widespread historical use in medicine and agriculture, and colistin due to its importance as an antibiotic of last resort. Migratory birds, including species of goose, likely harbor microbiomes richer in antimicrobial resistant bacteria compared to other microbiomes (34, 51). In one study, 50% of migratory birds encoded the emerging colistin resistance gene *mcr-1* within their microbiomes, while the most prevalent antimicrobial resistance genes in these microbiomes are against tetracycline (51). The dominant tetracycline resistance mechanism picked up by our experiment is drug efflux. As illustrated in the representative insert gene maps for this selection (**Figure 8A**), many tetracycline efflux pumps from this selection are syntenic to potential mobilization elements, including transposases, and phage- and plasmid-associated genes. In contrast, the dominant colistin resistance mechanism appears to be lipid A modification and our functional metagenomic library selections identified a potentially novel MCR enzyme homology, further confirming the presence of these concerning genes in migratory birds.

In summary, our experiments developing and testing METa assembly highlight several advantages of the method. First, the use of transposases to fragment metagenomic DNA has several benefits. It removes the need for costly capital equipment, such as sonicators, while providing the benefits of greater control and experimental flexibility seen with restriction enzyme-mediated fragmentation without the downsides of needing restriction site frequency or methylation dependence (77). Like sonication, tagmentation shows very little to no sequence bias, making it essentially random (40), but unlike sonication it can be applied to low biomass samples. The compatibility with low biomass samples is especially useful, as low biomass or rare microbiomes have forced other research groups being to pool independent samples (21, 35) or use potentially biased DNA amplification techniques (78, 79) to increase input DNA mass. One particularly relevant low biomass sample for METa assembly could be clinical swabs, which have been found on average to yield 371 ng of metagenomic DNA (80) which is well above the inputs we have used here (200 ng to 300 ng).

Second, the addition of mosaic end sequences to the DNA inserts allows the use of modern assembly cloning methods. Random fragmentation of metagenomic DNA by sonication or enzymatic digest (by restriction enzyme or fragmentases(67)) results in DNA fragments with little to no information about the DNA sequence at the fragment ends. This limits cloning of these fragments to lower efficiency methods like blunt ligation. More efficient assembly cloning protocols rely on incorporation of insert-matching sequences into the plasmid cloning site to allow insert-vector hybridization to drive ligation specificity and efficiency. As a result, current functional metagenomic library preparation methods have not been able to take advantage of high efficiency cloning techniques that have been available for more than a decade (44, 69). Assembly cloning and transposase fragmentation are therefore synergistic: without assembly cloning, tagmented DNA would be cloned *via* blunt ligation with no gains in efficiency, and without tagmentation, assembly cloning fails for lack of insert-vector hybridization.

Third, plasmids used in blunt end cloning, and therefore current functional metagenomic library preparation methods, must undergo phosphatase treatment to prevent self-ligation. This is not a problem in assembly cloning reactions due to the higher temperatures used. We omitted phosphatase treatment from our workflow and while this led to the detection of numerous empty vector colonies in our blunt ligation libraries, only a single METa assembly colony out of 115 tested was found to be lacking an insert (**Supplemental figures 3, 5, 7, 9, 10, and 11B**). We used inverse PCR to prepare linearized vectors for both blunt ligation and METa assembly which allowed us to incorporate mosaic end sequences into our vector of choice. This strategy is applicable to any plasmid that is readily amplified by PCR meaning that METa assembly can be easily incorporated into existing functional metagenomic library preparation workflows.

Finally, the METa assembly protocol presents significant time savings. The process of fragmenting DNA by tagmentation takes only 7 minutes followed by a 5-minute quench. In contrast, fragmentation by sonication with a *e.g.* a Covaris E220 instrument requires a 60 minute degas time on top of 20 minutes or more of fragmentation time. End repair with DNA polymerase to fill in 3’ overhangs following tagmentation takes 15 minutes (though it could likely be accomplished in less time) while end repair following physical fragmentation of DNA with an End-It kit requires 45 minute reactions and 10-minute heat inactivation. Most notably, assembly based cloning takes 15 minutes while it is generally recommended that blunt ended ligation reactions be allowed to react overnight for optimal efficiency. It has been suggested that functional metagenomic library preparation could be used in a rapid workflow for clinical detection of resistance genes (25). The time savings found in METa assembly, most notable in the cloning step, could be invaluable in such a workflow.

In conclusion, the synergistic combination of fragmentation by transposase and cloning by assembly allows METa assembly to prepare larger, less DNA greedy, and more robust functional metagenomic libraries. The advantages of METa assembly of functional metagenomic libraries could allow these valuable chemical biology tools to be prepared from sources previously out of reach including those of low biomass (such as from the built environment), requiring fast turnaround (such as in the clinic), or of limited availability (such as exotic or historical samples).

## AVAILABILITY

Sequencing data and scripts for data analysis and statistics are available at the Hartmann lab github: https://github.com/hartmann-lab

## AUTHOR CONTRIBUTIONS

**TSC**: Conceptualization, formal analysis, investigation, methodology, project administration, resources, supervision, validation, visualization, writing – original draft, writing – review and editing. **AGM**: Data curation, formal analysis, investigation, methodology, software, validation, visualization, writing – original draft, writing – review and editing. **EMH**: Funding acquisition, project administration, resources, supervision, writing – review and editing.

## ACKNOWLEDGEMENT

We would like to thank Dr. Rickard Sandberg for their deposition of the pTXB1-Tn5 plasmid in the Addgene depository. We wish to acknowledge and thank Dr. Neil L. Kelleher for sharing laboratory space and research support with TSC. We also wish to thank Dr. Gautam Dantas for critical reading of the manuscript.

## SUPPLEMENTAL TABLES AND FIGURES

**Supplemental table 1.**
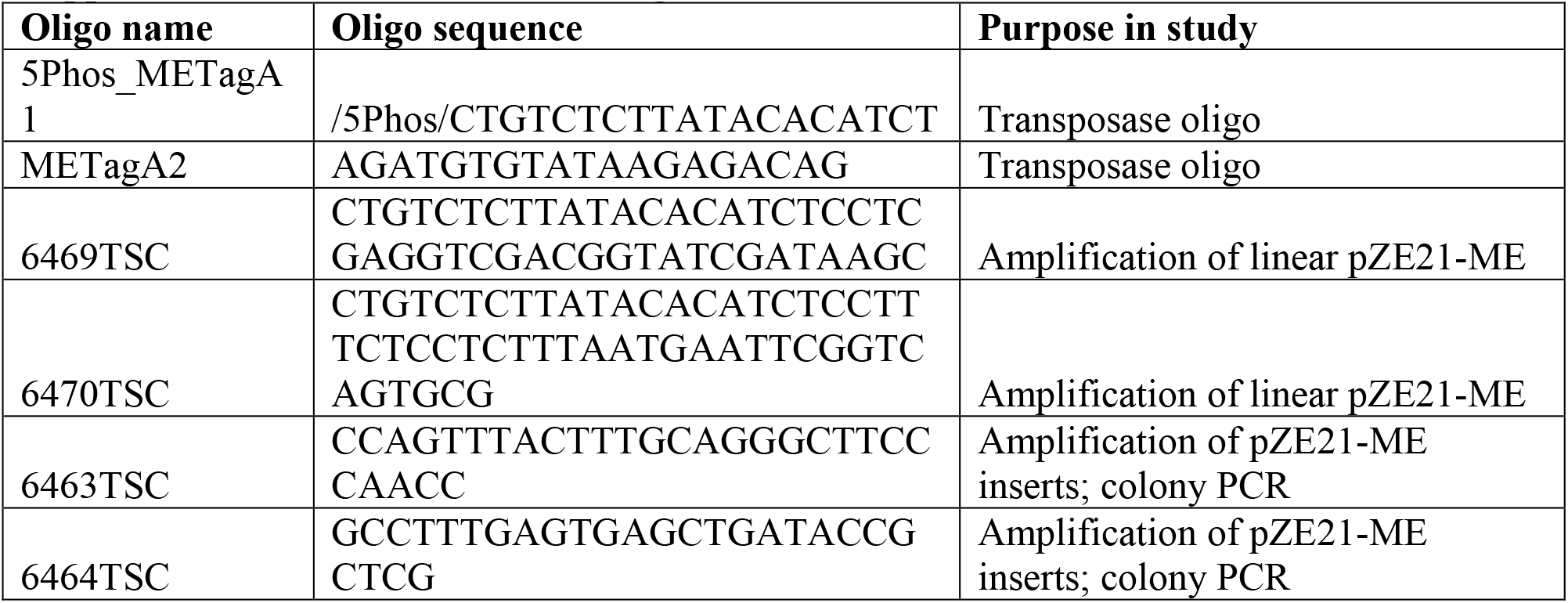
Primers and oligonucleotides used.

**Supplemental figure 1.**
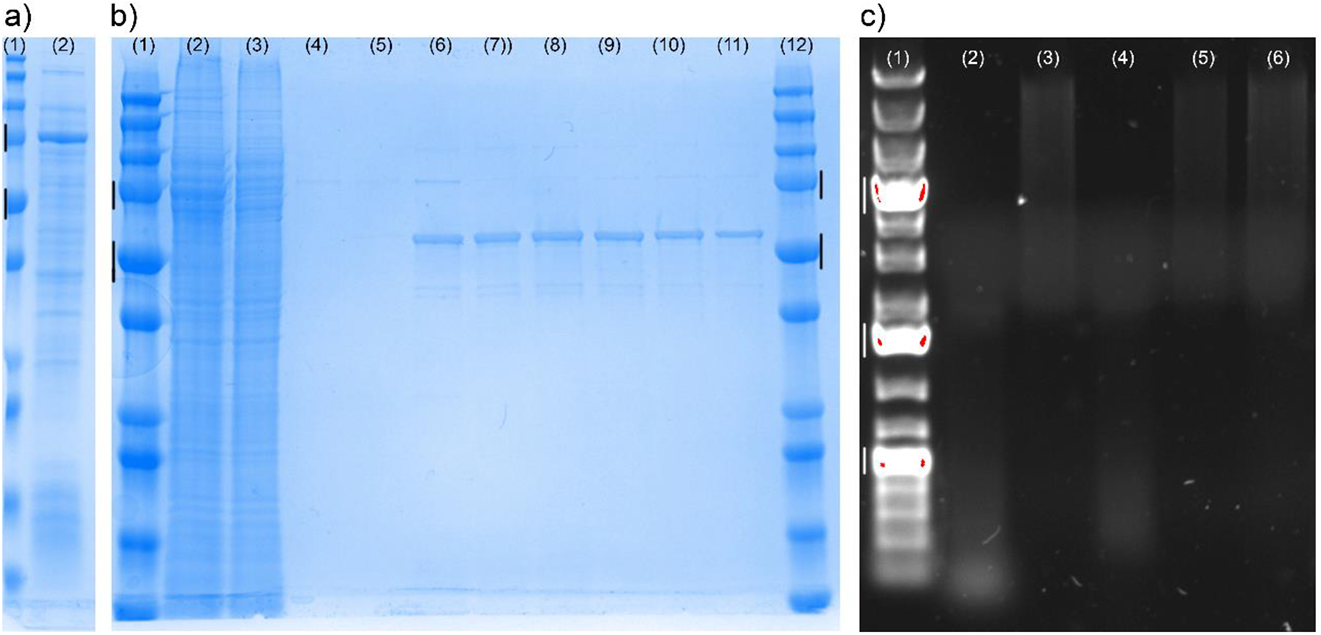
Expression, purification, and testing of transposase activity. a) Testing auto-induction expression of transposase. (1) Protein ladder with black bars next to 75 kDa and 50 kDa markers, (2) lysate from test expression culture. b) Transposase purification. (1) Protein ladder with 75 kDa and 50 kDa standards marked, (2) clarified lysate, (3) flow-through, (4) wash 1, (5) wash 2, (6) 24 hr test elution, (7) 48 hr elution, (8)-(11) elution washes 1 through 4, (12) ladder. Tn5-CDB fusion protein expected mass is ∼75 kDa, Tn5 post-cleavage expected mass is ∼50 kDa. c) Testing transposase activity. (1) DNA ladder with 5,000 bp, 1,500 bp, and 500 bp bands marked, (2) DNA treated with full transposase reaction, (3) DNA with transposase reaction minus enzyme, (4)/(5) same as (2)/(3) with column-based kit clean-up, (6) input metagenomic DNA without treatment.

**Supplemental figure 2.**
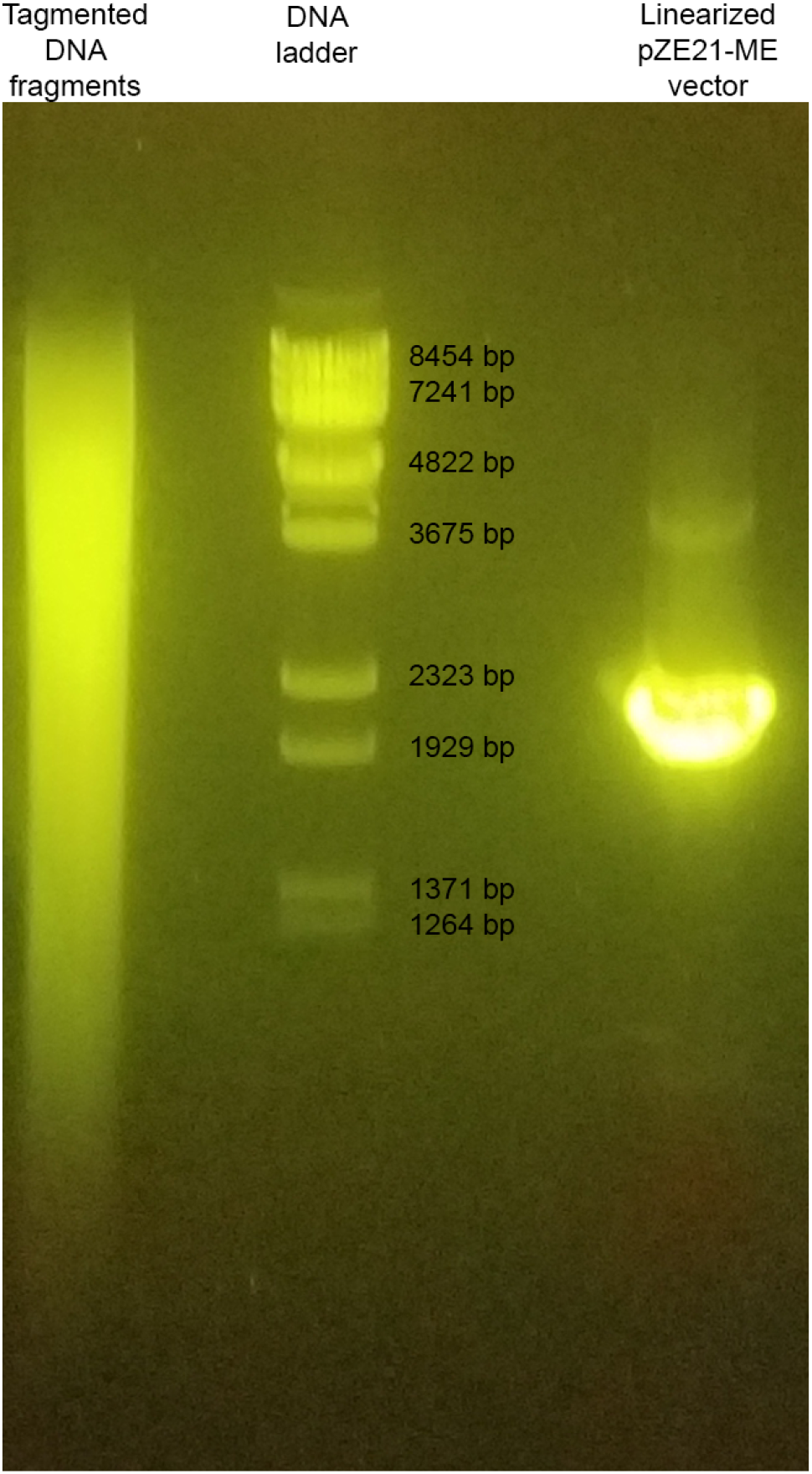
Size selection and purification of soil metagenomic tagmented DNA and inverse PCR prepared linear pZE21-ME vector. Tagmented soil metagenomic DNA (left of ladder) and inverse PCR amplified pZE21-ME (right of ladder) were purified from agarose gel. DNA fragments were excised from ∼1264 bp to ∼8454 bp and the major pZE21-ME band at ∼2280 bp was excised.

**Supplemental figure 3.**
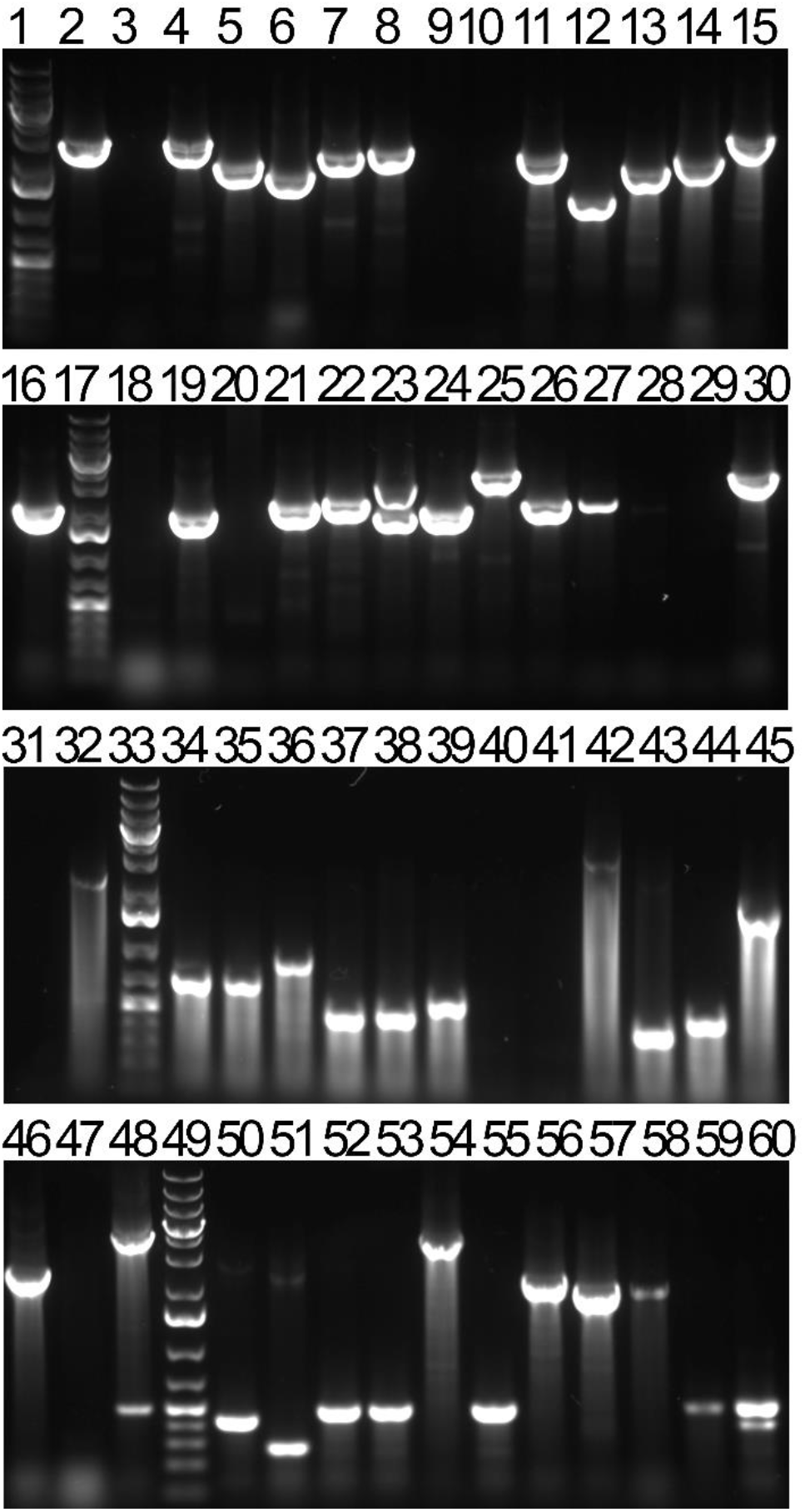
Colony PCR to determine library size and average insert length of soil metagenomic DNA prepared by METa assembly (In-Fusion and NEBuilder HiFi) or blunt cloning. Lanes 1, 17, 33, and 49: DNA ladders with brightest bands corresponding to 5 kb, 1.5 kb, and 500 bp. Lane 2: Single clone resulting from In-Fusion mediated METa assembly. Lanes 3-11, 12-21, and 22-31: Colonies from three replicate METa assemblies using NEBuilder HiFi. Lanes 32-41, 42-51, and 52-60: Colonies from triplicate blunt ligation cloning reactions. Lanes with bands at 500 bp correspond to amplification of vector backbone only (no inserts) while lanes with no band indicate colony PCR reaction failure

**Supplemental figure 4.**
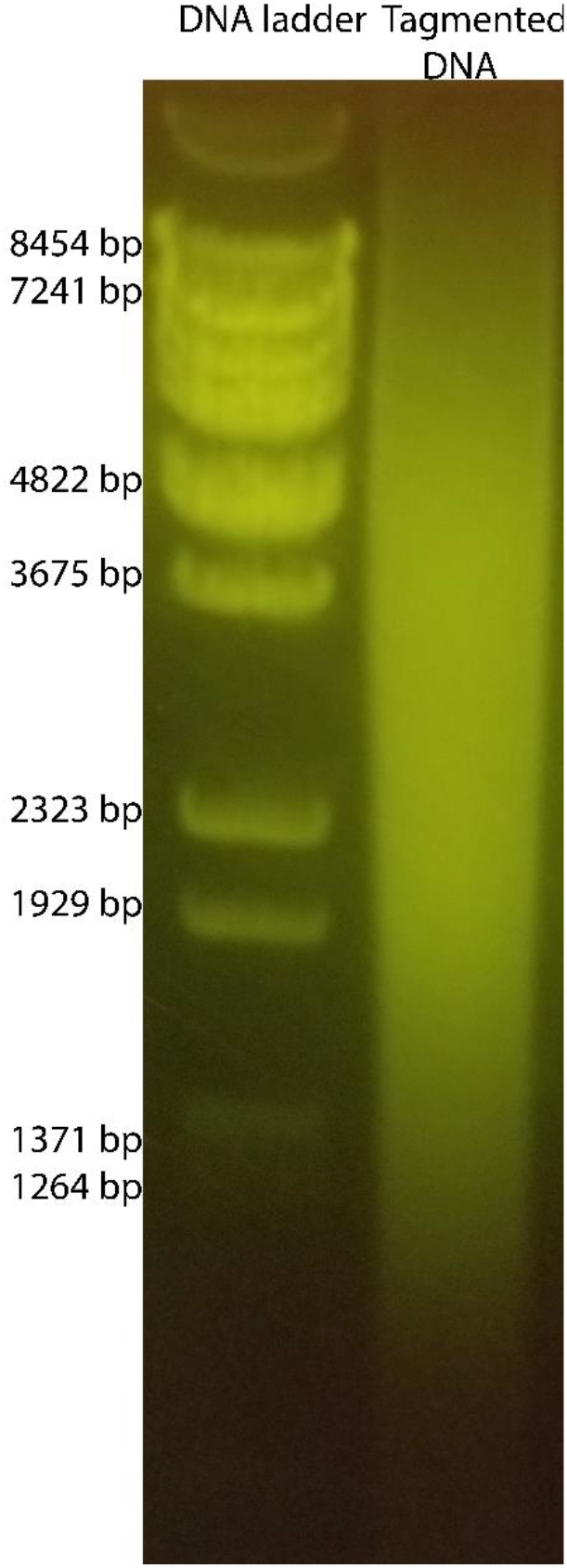
Size selection of mixed genomic tagmented DNA. Tagmented mixed ABC07, ABC10 genomic DNA was purified from agarose gel. DNA fragments were excised from ∼1264 bp to ∼8454 bp

**Supplemental figure 5.**
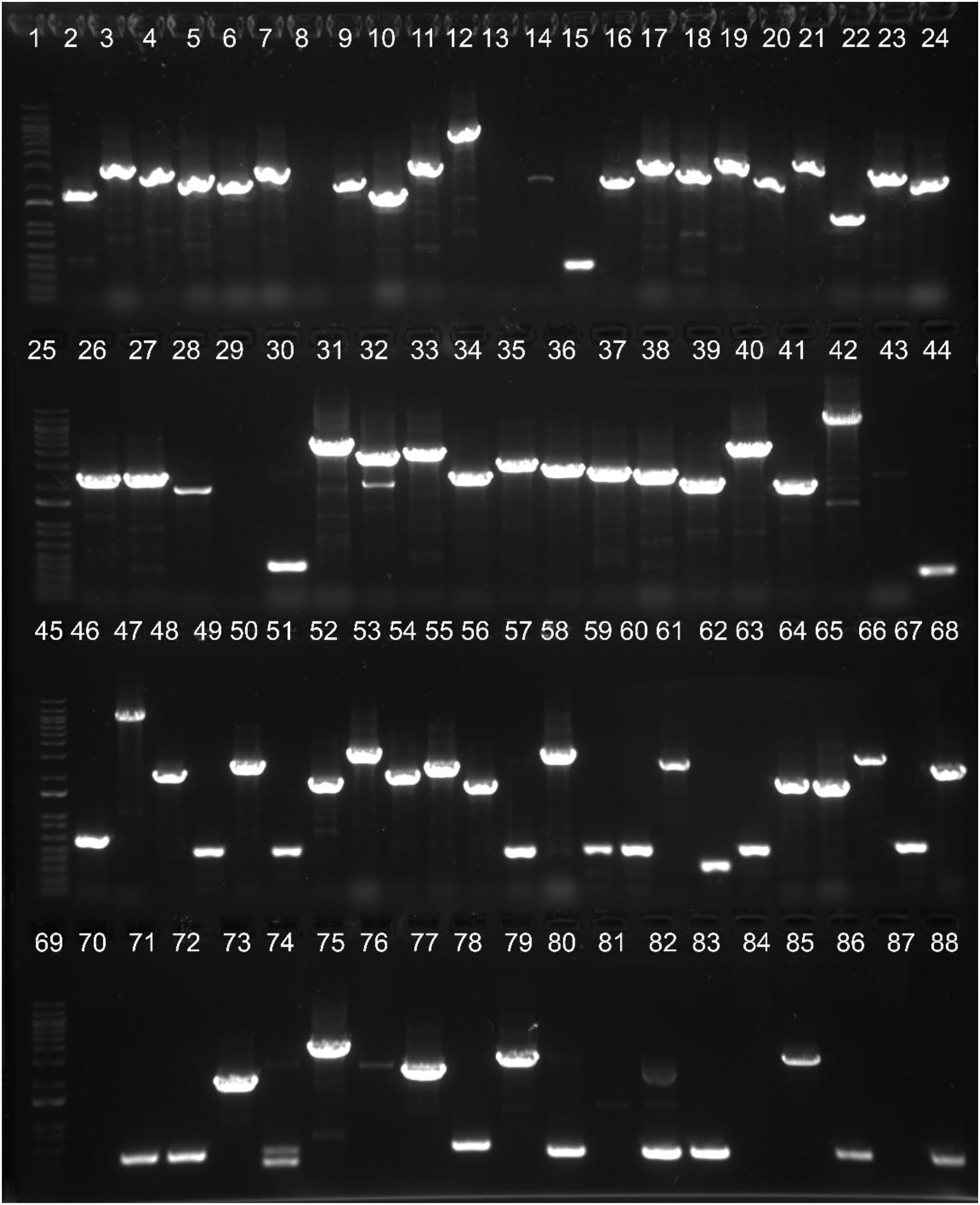
Colony PCR to determine library size and average insert length of mixed genome library prepared by METa assembly or blunt cloning. Lanes 1, 25, 45, and 69: DNA ladders with top and brightest bands corresponding to 20 kb and 1.5 kb, respectively. Lanes 2-24, 26-44: Colonies from three replicate METa assemblies using NEBuilder HiFi (13 colonies per replicate, lanes 15, 30, and 44 from negative control sham colonies). Lanes 46-68, 70-88: Colonies from triplicate blunt ligation cloning reactions (13 colonies per replicate, lanes 59, 74, and 88 from negative control sham colonies). Reactions resulting in 500 bp amplicons indicate carriage of a no inserts vector. Reactions resulting in no visible band indicate technical failure of the colony PCR reaction.

**Supplemental figure 6.**
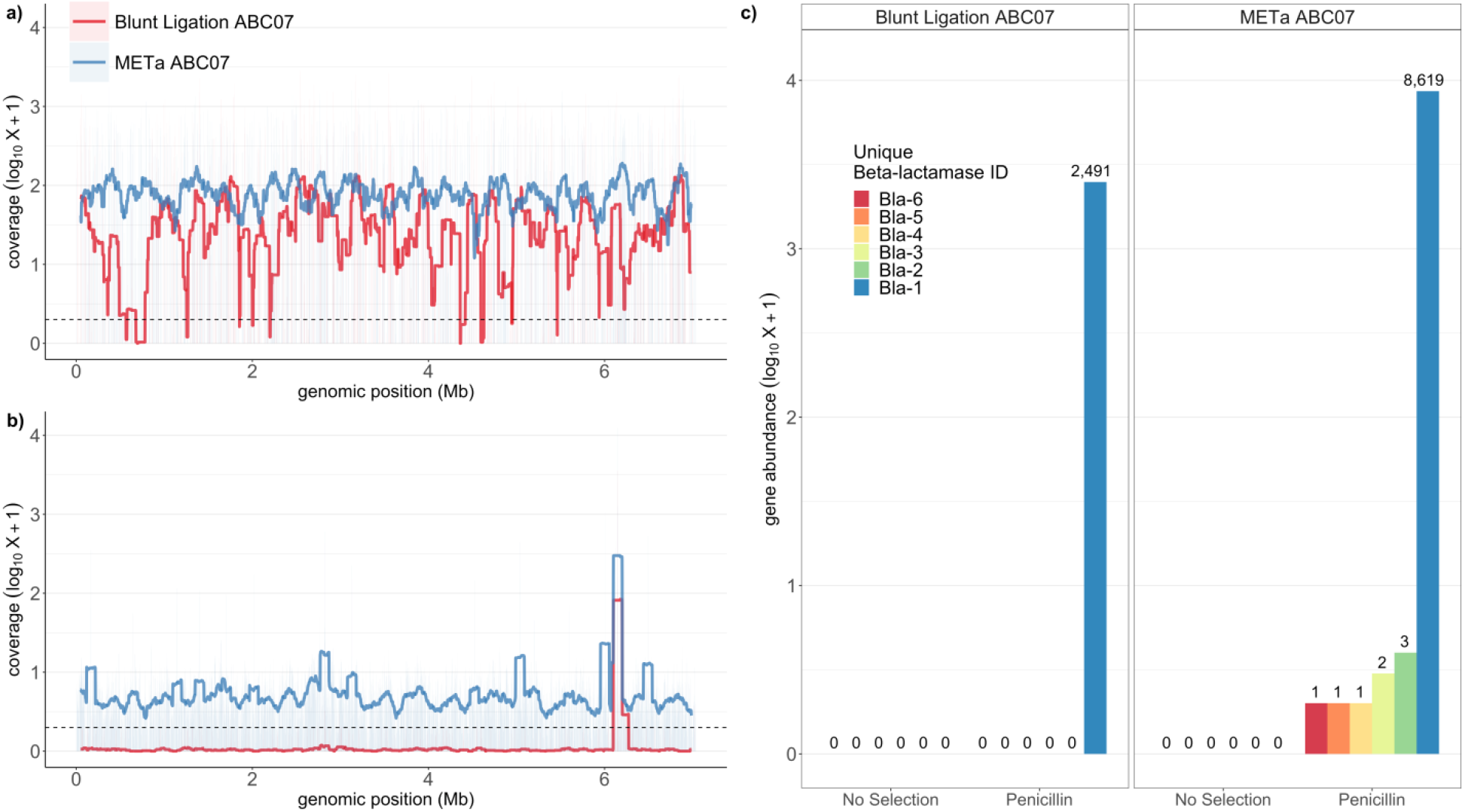
Assembly and blunt ligation library coverage of ABC07 genome with and without penicillin selection. a) Nucleotide depth of coverage for ABC07 genome by functional metagenomic library prepared by assembly (blue) or blunt ligation (red). Coverage is smoothed to a 1 kb resolution. b) Same as in a) but sequenced libraries were first subjected to selection on agar plates containing 1 mg/ml penicillin. c) Gene abundance in post-penicillin selection reads for each of six predicted ABC07 β-lactamase genes that had read number >0. From Bla-1 to Bla-6 respective NCBI accession numbers are WP_087694996.1, WP_003211977.1, WP_003216184.1, WP_087694154.1, WP_008436310.1, WP_003207471.1.

**Supplemental figure 7.**
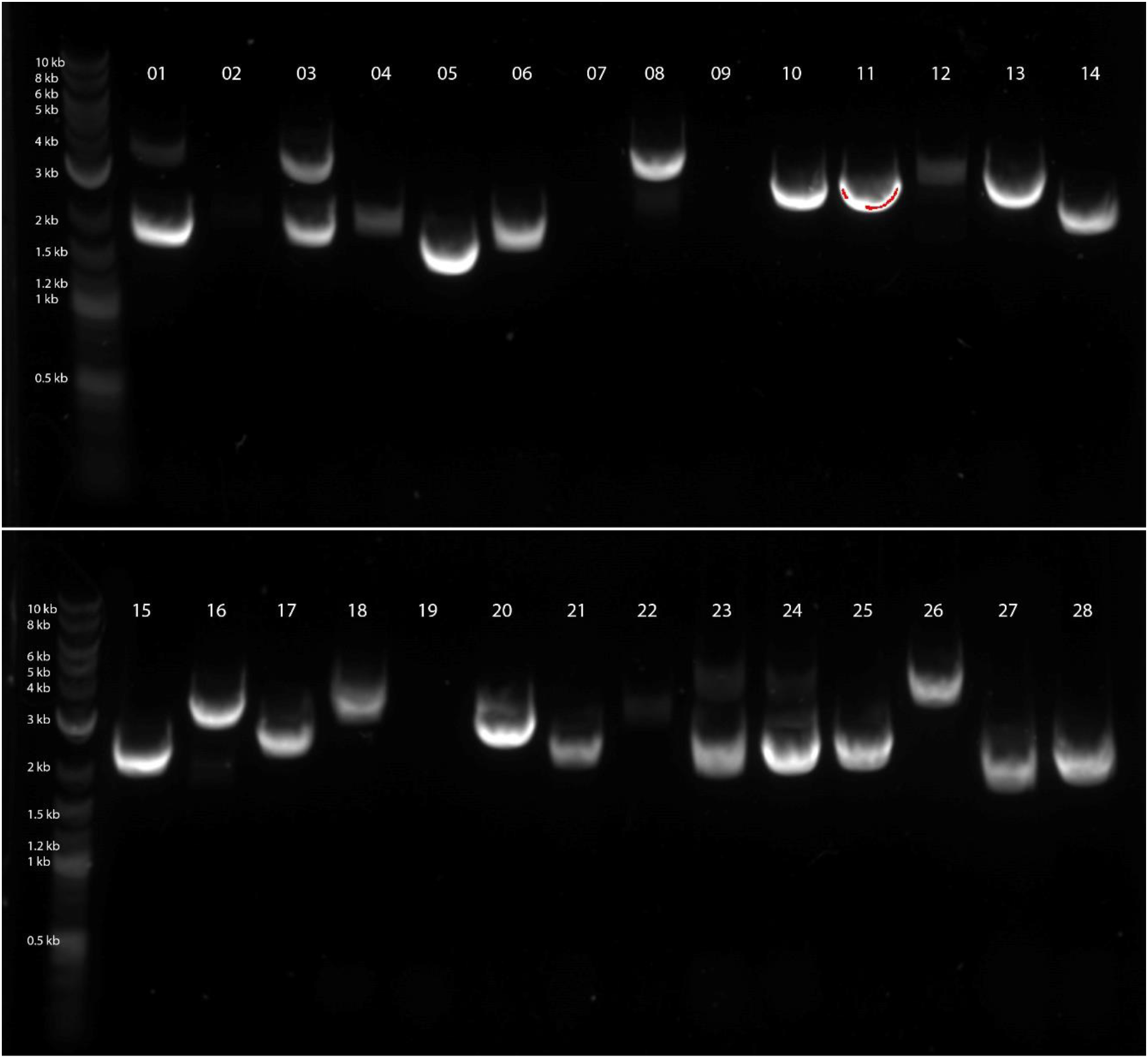
Representative colony PCR to determine average insert and library size of 5 μg soil library. The first lane of each gel contains ladder, numbered lanes correspond to 28 colony PCR reactions. Reactions for colonies 2, 7, 9, and 19 showed no amplicon indicating reaction failure. Lack of reactions producing 500 bp amplicons indicates all successful reactions contained an insert.

**Supplemental figure 8.**
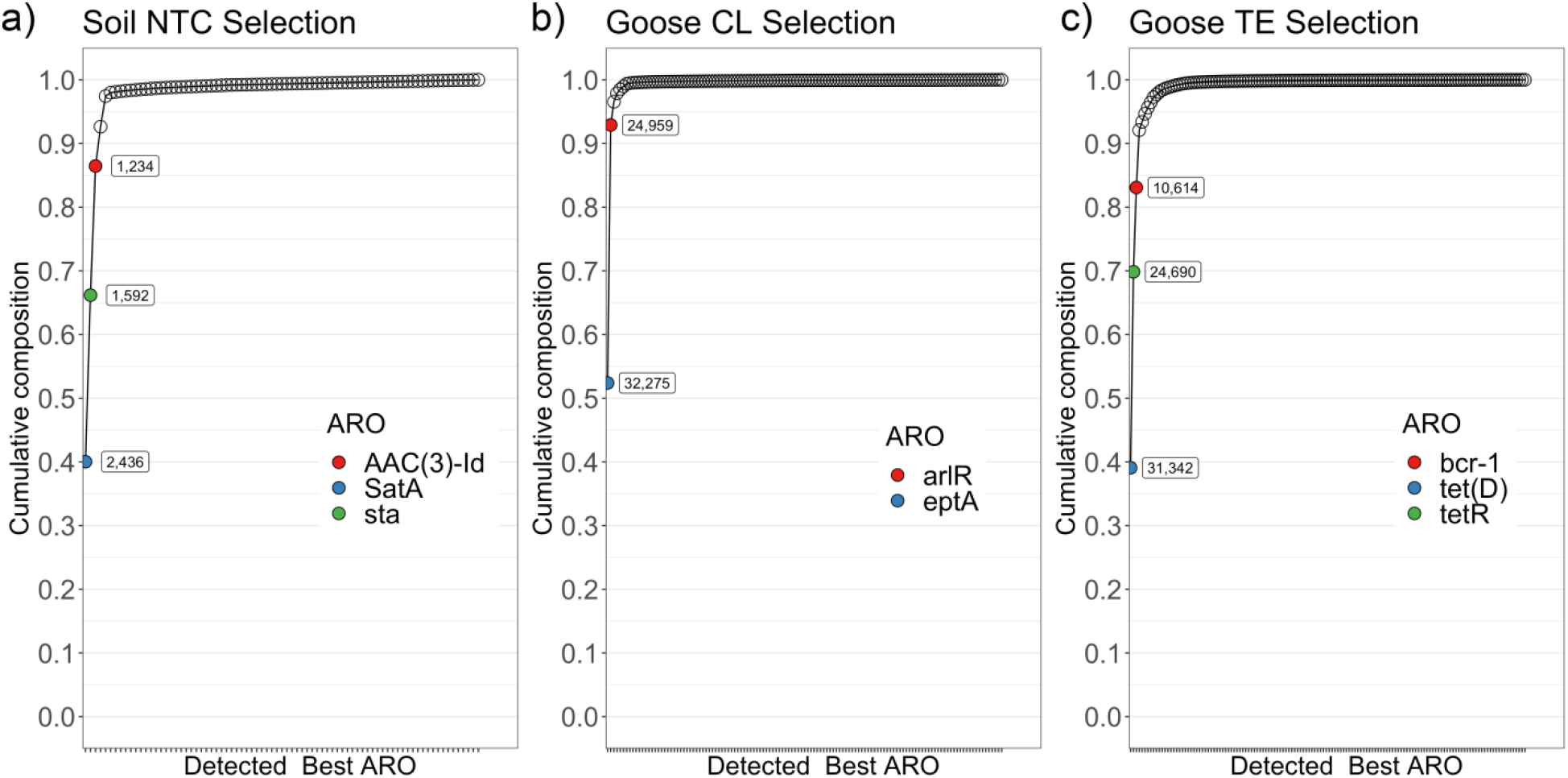
Antibiotic resistance gene family abundance. Major gene families predicted by CARD Antibiotic Resistance Ontology for a) soil microbiome selected on nourseothricin (NT) or goose gut microbiome selected on b) colistin (CL) or c) tetracycline (TE).

**Supplemental figure 9.**
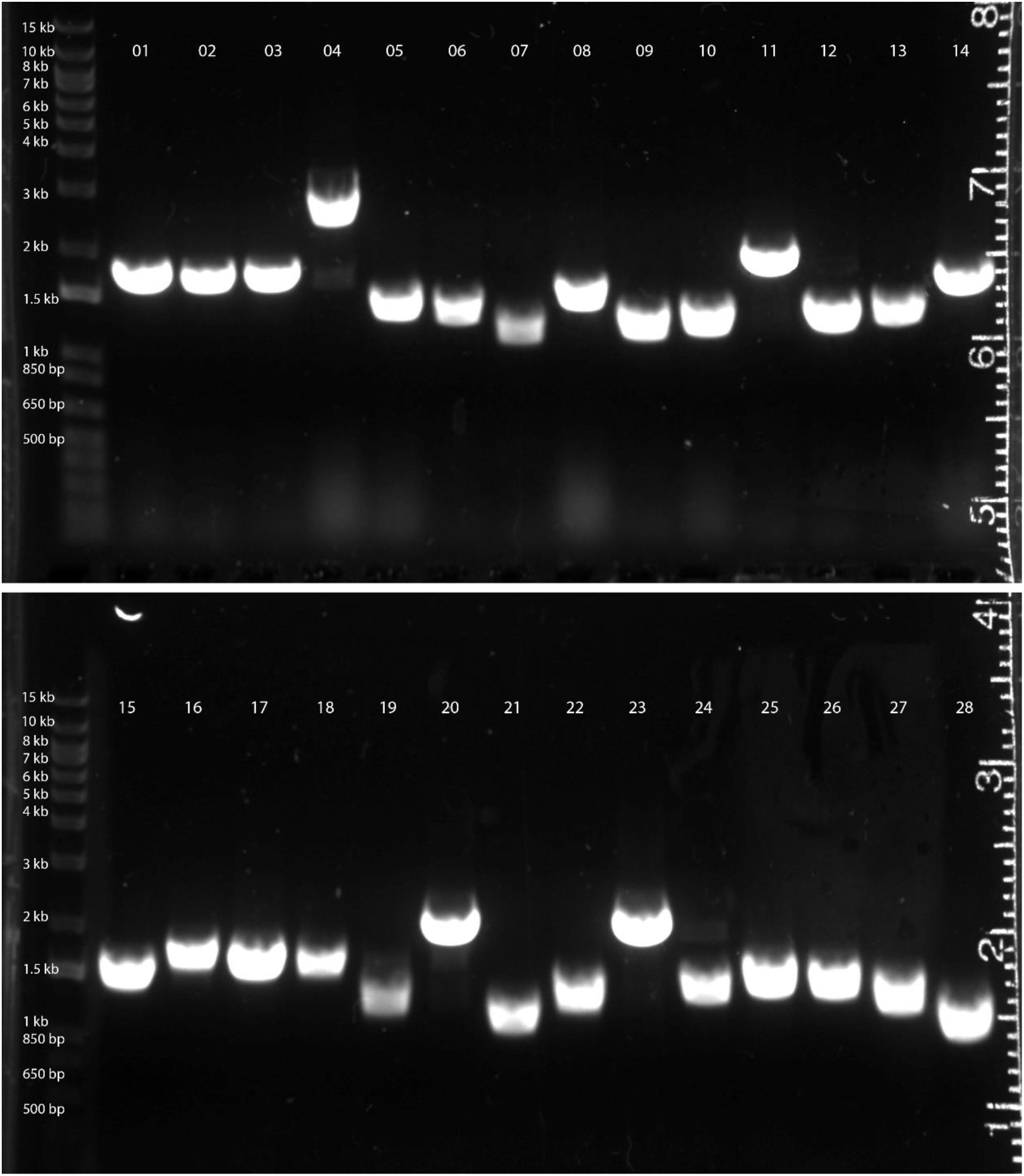
Colony PCR to determine average insert size and library size of 10 kb amplicon test library. The first lane of each top and bottom gel contain ladder, numbered lanes correspond to 28 colony PCR reactions. All tested colonies contained plasmids with inserts as indicated by all amplicons exceeding 500 bp in size.

**Supplemental figure 10.**
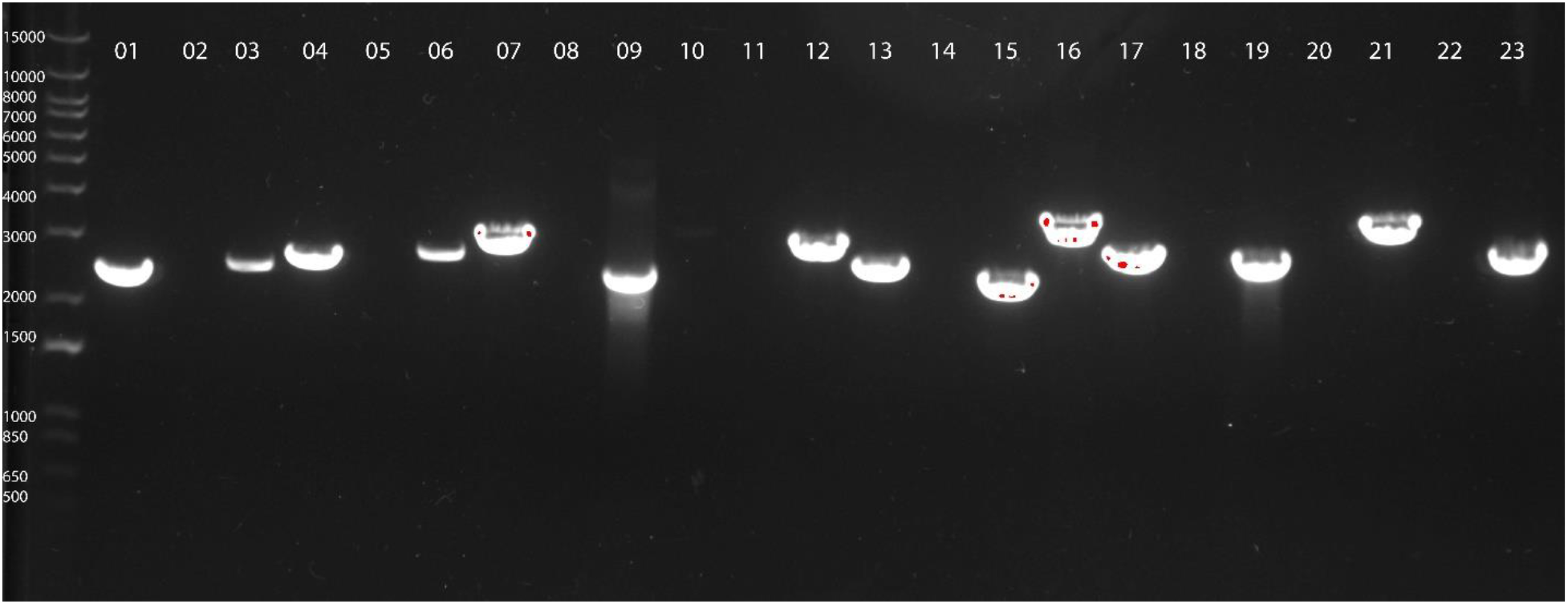
Colony PCR to determine average insert size and library size of 250 ng soil functional metagenomic library. The first lane contains ladder, numbered lanes correspond to 24 colony PCR reactions. Reactions 2, 5, 8, 10, 11, 14, 18, 20, and 22 failed. All successful amplification reactions contained plasmids with inserts as indicated by all amplicons exceeding 500 bp in size.

**Supplemental figure 11.**
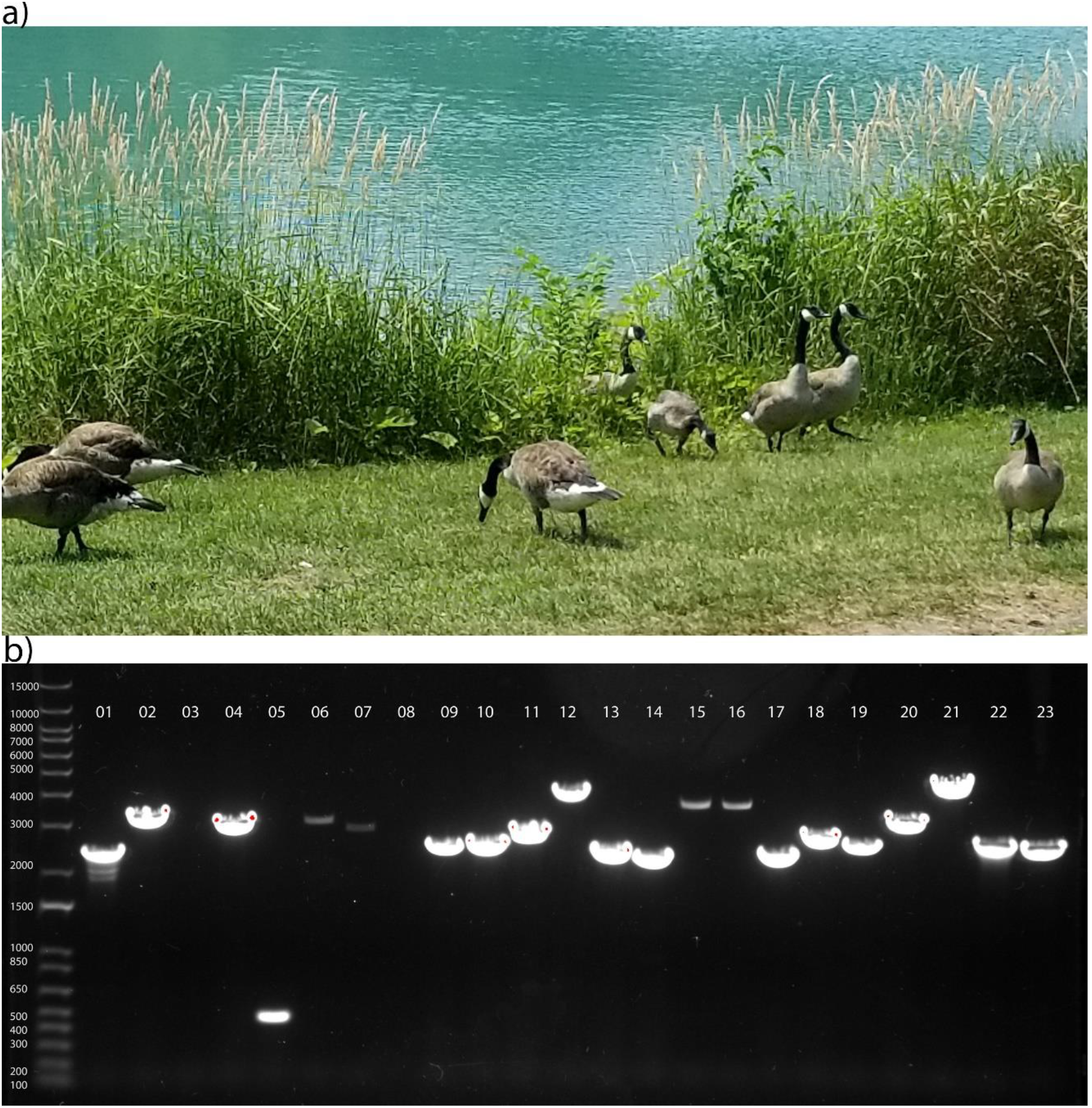
Low input goose fecal microbiome functional metagenomic library. a) The gaggle of geese putatively identified as *Branta canadensis* from which the fecal pellet was obtained. b) Colony PCR to identify average insert size and estimate total library size. The first lane contains ladder and the remaining lanes contain PCR reactions performed on 23 library colonies. Reactions 3 and 8 failed while the 500 bp amplicon in reaction 5 corresponds to 1/21 colonies containing no insert.

**Supplemental figure 12.**
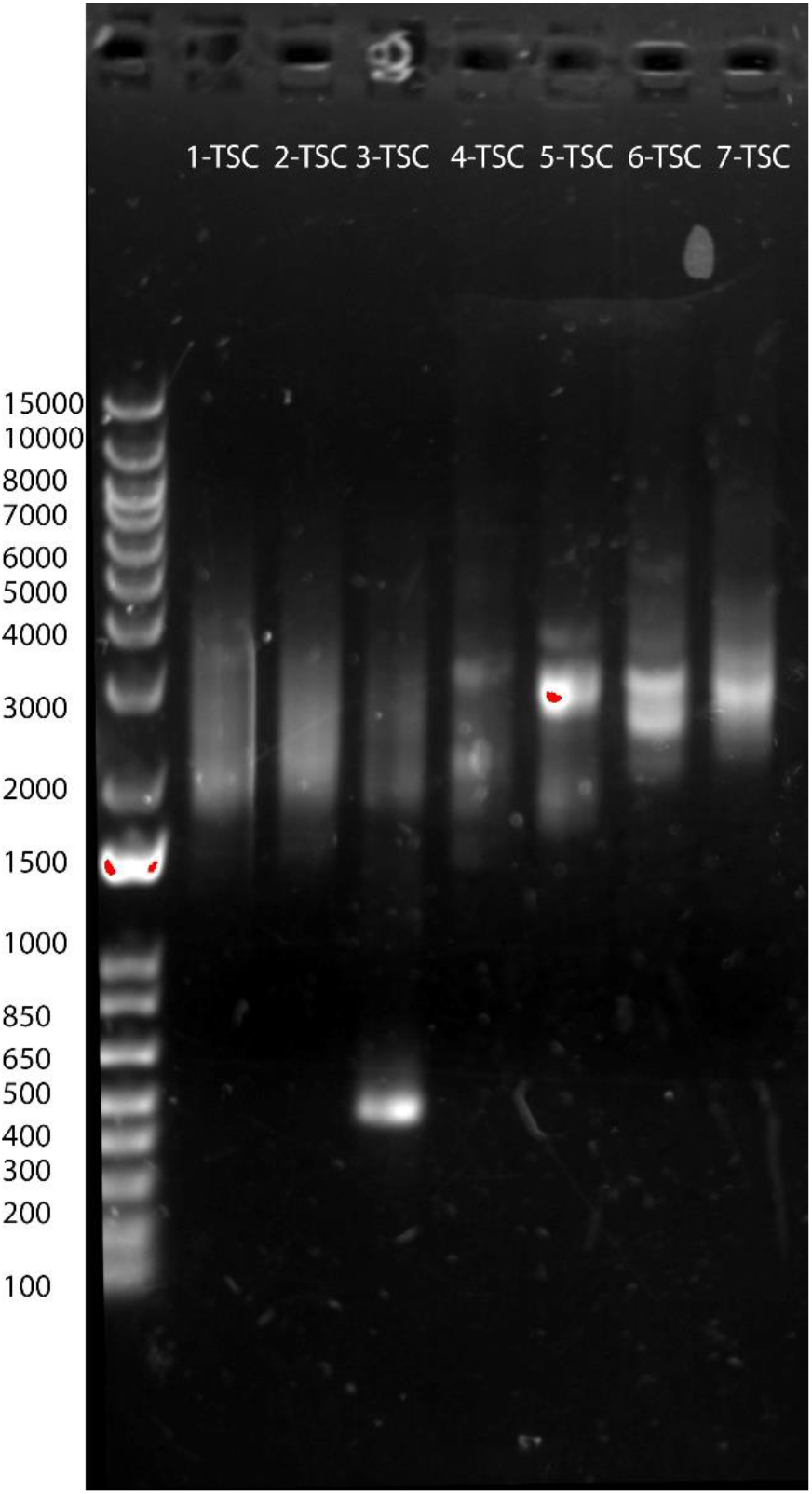
Amplicons from functional metagenomic library minipreps for sequencing. From left to right: Ladder, (1-TSC) pooled triplicate ABC07/10 library prepared by METa assembly, no antibiotics, (2-TSC) same as 1-TSC following selection on 1 mg/ml penicillin, (3-TSC) pooled triplicate ABC07/10 library prepared by blunt ligation, no antibiotics, (4-TSC) same as 3-TSC following selection on 1 mg/ml penicillin, (5-TSC) 5 μg soil microbiome library following selection on 64 μg/ml nourseothricin, (6-TSC) 300 ng goose fecal microbiome library following selection on 8 μg/ml tetracycline, (7-TSC) same as 6-TSC but selection on 4 μg/ml colistin.

**Supplemental figure 13.**
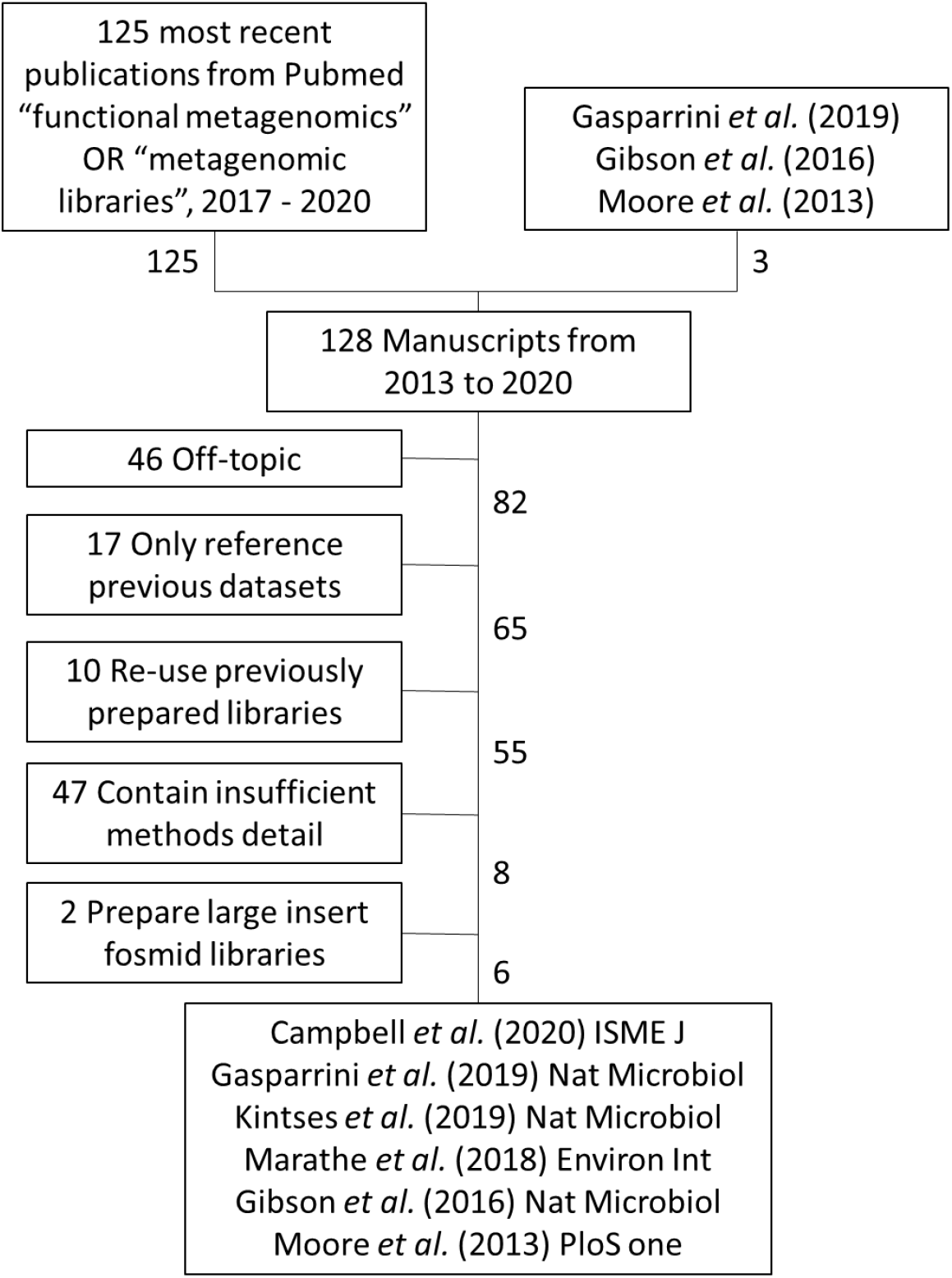
Literature search for functional metagenomic library preparation details. See methods for details.

